# Mathematical modeling suggests cooperation of plant-infecting viruses

**DOI:** 10.1101/2021.12.17.473153

**Authors:** Joshua Miller, Tessa M. Burch-Smith, Vitaly V. Ganusov

## Abstract

Viruses are major pathogens of agricultural crops. Viral infections often start after the virus enters the outer layer of a tissue or surface and many successful viruses, after local replication in the infected tissue, are able to spread systemically. Quantitative details of virus dynamics in plants, however, have been poorly understood, in part, because of the lack of experimental methods allowing to accurately measure the degree of infection in individual plant tissues. Recently, by using flow cytometry and two different flourescently-labeled strains of the Tobacco etch virus (TEV), Venus and BFP, kinetics of viral infection of individual cells in leaves of *Nicotiana tabacum* plants was followed over time [1]. A simple mathematical model, assuming that viral spread occurs from lower to upper leaves, was fitted to these data. While the the original model could accurately describe the kinetics of viral spread locally and systemically, we also found that many alternative versions of the model, for example, if viral spread starts at upper leaves and progresses to lower leaves or when virus dissemination is stopped due to an immune response, provided fits of the data with reasonable quality, and yet with different parameter estimates. These results strongly suggest that experimental measurements of the virus infection in individual leaves may not be sufficient to identify the pathways of viral dissemination between different leaves and reasons for viral control; we propose experiments that may allow discrimination between the alternatives. By analyzing the kinetics of coinfection of individual cells by Venus and BFP strains of TEV we found a strong deviation from the random infection model, suggesting cooperation between the two strains when infecting plant cells. Importantly, we showed that many mathematical models on the kinetics of coinfection of cells with two strains could not adequately describe the data, and the best fit model needed to assume i) different susceptibility of uninfected cells to infection by two viruses locally in the leaf vs. systemically from other leaves, and ii) decrease in the infection rate depending on the fraction of uninfected cells which could be due to a systemic immune response. Our results thus demonstrate the difficulty in reaching definite conclusions from extensive and yet limited experimental data and provide evidence of potential cooperation between different viral variants infecting individual cells in plants.

## Introduction

With a burgeoning population expected to reach 9.7 billion by the midpoint of the 21st century, humans’ lifeblood, food and water, will be evermore difficult to protect and sustain over time [2]. Under these circumstances, dependence on agriculture will only increase [3]. Food crops, however, are vulnerable to numerous entities, including but not limited to animal pests, fungi, bacteria, and viruses. Viral infections especially can devastate food-crops, with documentation of such infections being identified as early as eighth-century Japan; however it was not until the nineteenth century that it was known and accepted that microscopic agents like viruses could cause diseases in plants [4, 5].

Mathematical models have been widely used to understand virus-plant interactions. For example, early studies investigated how the virus concentration in the inoculum influences the number of lesions formed by the virus on plant leaves [6–9]. More recent studies investigated virus dynamics in individual plant cells or in the whole plant [1, 10–12]. Most studies, however, focused on understanding epidemiological spread of viral disease in plant populations with the aim of controlling viral spread and limit damage to agricultural crops [13–23].

Mechanisms of viral spread within individual plants remain incompletely understood. Usually infection of a single cell or a small group of cells occurs via mechanical means or by an animal or insect vector. After replication in the inoculation site, virus moves to neighboring cells through plasmodesmata – pores between individual cells in the leaf [24]. The replication-movement process is repeated until the virus enters the vasculature. It has been experimentally demonstrated that viral distribution via the vasculature follows the path of sugar distribution, i.e., from source to sink tissues, with a strong sink like roots receiving a larger portion of the viral cargo. Once arriving at sink tissue, the virus exits via the plasmodesmata and enters neighboring cells. From there viruses use plasmodesmata once again to invade the ground tissue [25, 26]. In some cases, however, viruses can be introduced directly into the vasculature resulting in rapid infection of sink leaves.

Different methods have been used to measure the infection degree of a given leaf in the plant including ELISA for viral proteins and PCR for viral genomes [11, 27]. However, these methods are semi-quantitative and typically do not allow to measure degree of infection of individual cells in the leaf. Recently, a new method to measure the frequency of infection of cells in plant leaves with the use of flow cytometry was developed [1]. In their experiments, Tromas *et al*. [1] infected lower (3^rd^) leaves of 4 week old *Nicotiana tabacum* plants with two strains of Tobacco etch virus (TEV), TEV-Venus and TEV-BFP, carrying different fluorescent proteins. At different times after the infection cells (protoplasts) were isolated from individual leaves, and the fraction of protoplasts infected with either or both viral variants was quantified using flow cytometry [1]. Flow cytometry allowed the measurement of virus infection in thousands of individual cells, thus, providing exclusive quantitative information about kinetics of TEV infection in *N. tabacum*.

Tromas *et al*. [1] performed several important analyses including calculation of basic reproductive number and multiplicity of infection of cells by different viruses. In addition, the authors developed a detailed mathematical model of how the virus spreads over time from the 3^rd^ leaf to other leaves and fitted the model to experimental data. Importantly, the model was able to accurately describe virus dissemination and predicted that viral spread was similar within the leaves. One major difference between infection rates was due to different import rates of the virus from the lower to upper leaves [1].

Here we built upon this pioneering work and further analyzed experimental data of Tromas *et al*. [1] with use of mathematical models. Because exact pathways of TEV dissemination in *N. tabacum* plants have not been firmly established and may depend on the age of plants and details of virus inoculation, we therefore investigated whether alternative mathematical models of TEV dissemination may be consistent with the data. Surprisingly, we found that indeed many different routes of TEV dissemination (e.g., when the initial infection first spreads in top (7^th^) leaf and then disseminates to lower leaves) are well consistent with experimental data, even though some of such models fitted the data with slightly reduced quality (as evaluated by AIC, [28]). By analyzing kinetics of coinfection of individual cells by two TEV variants we found that coinfection does not proceed randomly; rather, cells are more likely to be coinfected with two viruses than infected with either of the variants suggesting cooperativity in infection (or that plant cells vary in susceptibility to infection). Our results suggest that understanding pathways of virus dissemination in plants will be difficult using only data on virus infection in individual leaves and may likely require specific experiments.

## Materials & Methods

### Data

Specific details of how infection of plants had been performed are given in the previous publication [1]. In short, 4 week old N. *tabacum L. cv. Xanthi* plants, a widely used model plant host [29], were inoculated into the 3^rd^ leaf with equal mixture of two Tobacco etch virus (TEV) variants, TEV-BFP and TEV-Venus. These two viral strains express blue and yellow fluorescent proteins, respectively. Preliminary work demonstrated that expression of these proteins does not impair growth kinetics of the viral variants [1]. The plants were infected by infectious saps, applied using a rub-inoculation technique. To measure the kinetics of viral dissemination 3^rd^, 5^th^, 6^th^, and 7^th^ true leaves of individual plants were removed at days 3, 5, 7 and 10 post inoculation; five plants per time point were analyzed. Leaf 4 was skipped because it did not show any infection under the current experimental conditions. From these leaves, plant cells (protoplasts) were isolated and the number of protoplast expressing none, one, or both of the two fluorescent proteins was measured with flow cytometry. The data have been formatted and are available as a supplement to this paper.

### Mathematical models

#### Original virus dissemination model by Tromas *et al*. [1]

To predict kinetics of infection of the inoculated leaf and dissemination of infection to other leaves in the plant Tromas *et al*. [1] developed a novel mathematical model. The model tracks the fraction of infected cells in a *k*^th^ leaf, *I_k_* over time with *S_k_* being the fraction of susceptible cells. In the the model a cell infected with either of two viral variants or both viral variants is considered to be infected. The model assumes that infection starts at leaf 3 and then proceeds in the leaf *k* = 3 at a rate *β* and disseminates to upper leaves (leaves 5, 6, 7) at a rate proportional to the total infection rate of the leaves below a given leaf *k* at a rate *χ_k_* (Figure 1A). When the virus reaches other leaves, infection also proceeds locally at a rate *β*. Local virus dissemination at a *k*^th^ leaf stops when the fraction of infected cells reaches a critical level *ψ_k_*:

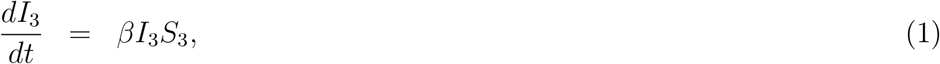

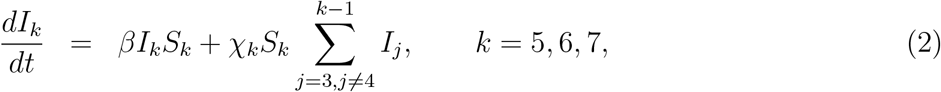

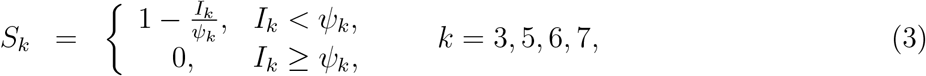

where *β* is the rate of infection of uninfected cells in the leaf during to local viral spread in the leaf, *χ_k_* is the rate of virus infection of other (upper) leaves, and *ψ_k_* is the level at which infection of new cells in leaf *k* stops (see eqns. (S.1)–(S.4) for full set of equations of the model). Initial conditions for the model are *I_k_*(0) = *I*_0_ if *k* = 3 and *I_k_*(0) = 0, otherwise, and *S_k_*(0) = 1. In total, this model has 9 parameters to be estimated from the data.

**Figure 1:**
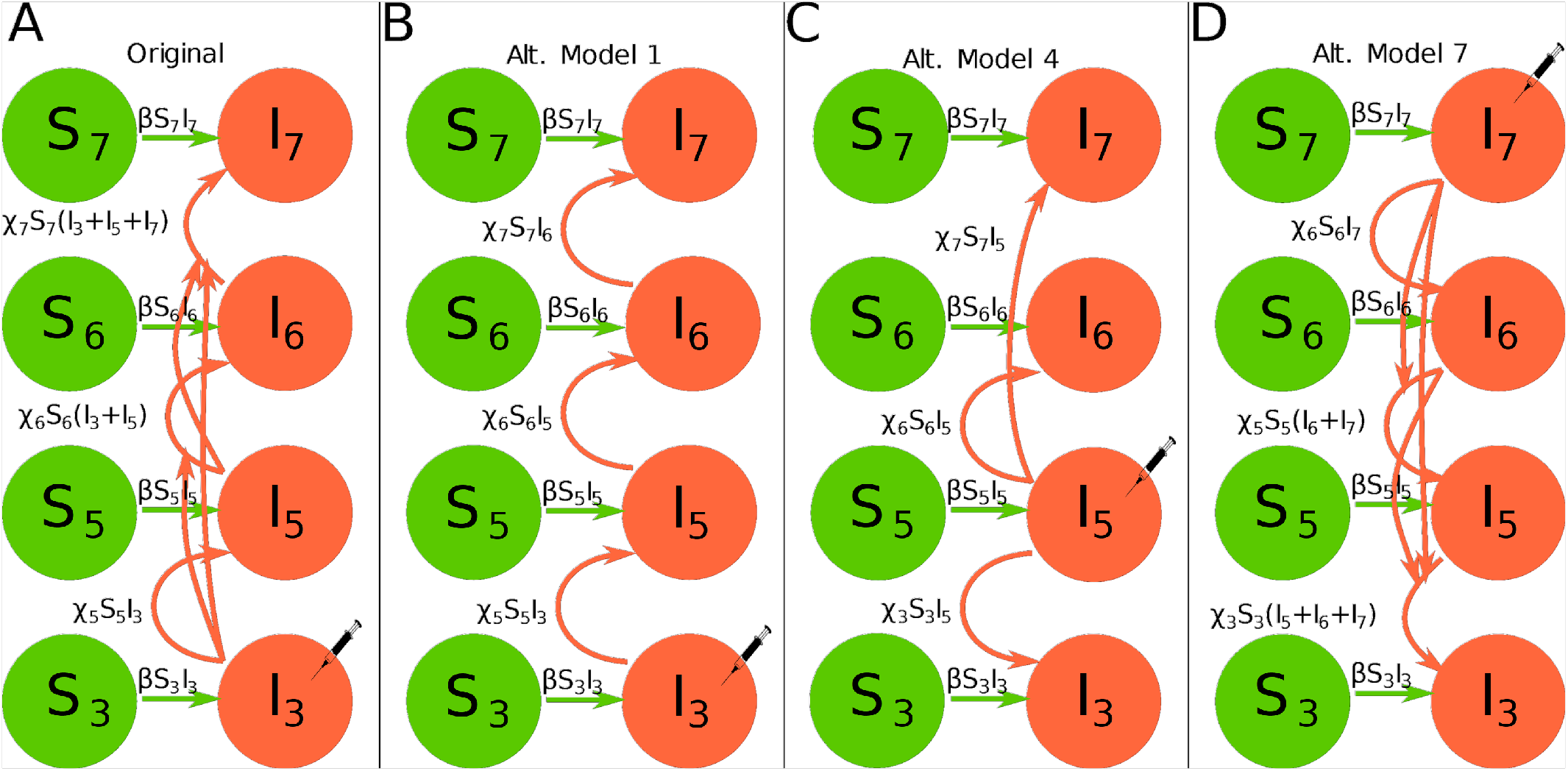
Examples of several mathematical models of virus spread in plants analyzed in this paper. In experiments of Tromas *et al*. [1], two different viruses (“Venus” and “BFP”) were rubbed into leaf 3 and the fraction of infected leaf cells (protoplasts) was followed by flow cytometry over time (see Materials and Methods for more detail). In schematics, *S_k_* and *I_k_* denote uninfected and infected cells in the *k*^th^ leaf, respectively, and syringe indicates the primary place where infection started in the model. Arrows denote the process of leaf infection (at a rate *β*) and transmission of infection between leaves (at a rate *χ*). In the original Tromas *et al*. [1] model (A, eqns. (1)–(3)), infection starts at leaf 3 and then transported to other leaf at a rate proportional to the total fraction of infected cells in leaves below. In alternative model 1, infection starts with leaf 3 but upper leaves are only infected by the leaves just below them (B, eqn. (4)). In alternative model 4, infection starts in leaf 5 and then processed to leaves above or below leaf 5 similar to the alternative model 1 (C, eqn. (11)). Finally, in the alternative model 7 infection starts at the upper leaf 7 and proceeds to leaves below similarly to the original Tromas *et al*. [1] model (D, eqn. (15)). Other alternative models are described in the Materials and methods.

**Figure 2:**
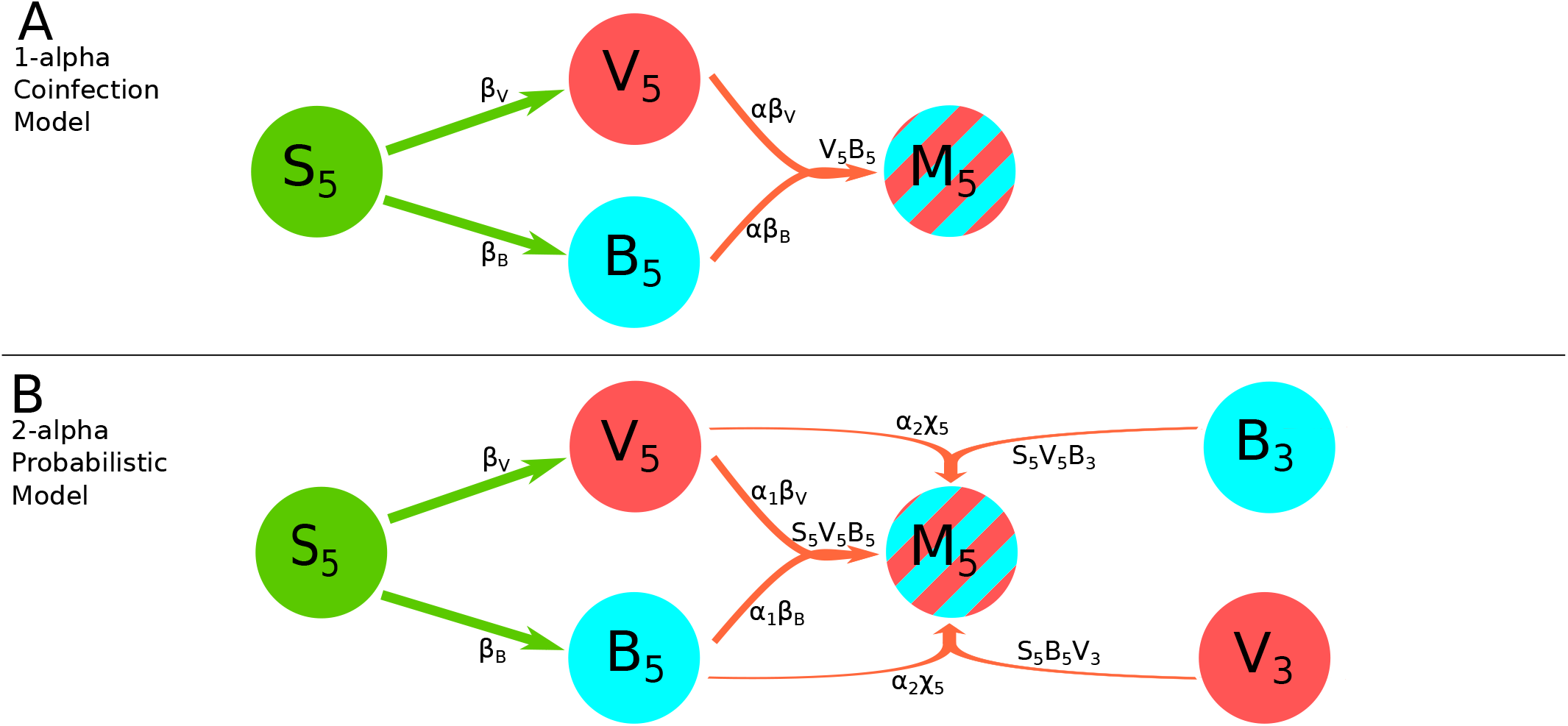
Examples of schematics of alternative mathematical models for coinfection with two viruses. These diagrams show the infection pathways of infection by TEV-Venus (*V_k_*) or TEV-BFP (*B_k_*) in the leaf 5 of the plants, and how these strains combine to form coinfected cells (*M_k_*) in the 1-alpha conifection model (A, eqn. (21)) or 2-alpha Probabilistic Model (B, eqn. (25)). Major model parameters such as *β* and *χ_k_* have the same meaning as in the previous models (e.g., Figure 1). We only show what happens in leaf 5 because in the 2-alpha Probabilistic Model (B, eqn. (22)), the connections between higher leaves become very complicated and difficult to illustrate in a figure such as this one. Like Figure 1, arrows represent the transmission of virions. In the 1-alpha Coinfection Model (A, eqn. (21)), coinfection comes from the combination of the Venus and BFP viruses within leaf 5 only. In the 2-alpha Probabilistic Model (B, eqn. (22)), coinfection comes also from the combination of Venus and BFP virions in leaf five, but is also fed by the combination of virions imported from leaf 3 and combining with their opposite, e.g. Venus from leaf 3 combining with BFP from leaf 5. Two separate alpha terms are used to distinguish dynamics between within-leaf growth and growth from virions imported from lower leaves.

#### Alternative virus dissemination models for the total leaf infection

While the original mathematical model of Tromas *et al*. [1] seems logical we sought to investigate whether alternative mathematical models of virus dynamics within individual leaves and virus dissemination to other leaves in the plant may be consistent with experimental data. In most of these alternative models we use the same nomenclature for the model parameters (*I*_0_, *β, χ_k_*, and *ψ_k_*) as in Tromas *et al*. [1].

- **Alternative model 1.** In this model the dynamics of infection of the leaf 3 is given in eqn. (1), and instead of summing the infection from all the leaves below, we suppose that only the leaf immediately below the one in question can infect it. The dynamics of uninfected leaves is given by eqn. (3). Dynamics of infection in other leaves is described by the following equations (see Figure 1B):

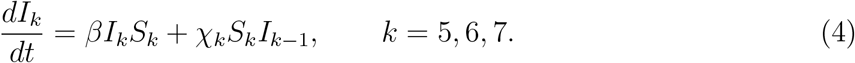 Initial conditions for the model are *I_k_*(0) = *I*_0_ if *k* = 3 and *I_k_*(0) = 0, otherwise, and *S_k_*(0) = 1.
- **Alternative Model 2.** Leaf 3 infects only leaf 5 which then infects leaves 6 and 7. Leaf 6 also contributes to the infection of leaf 7. Infection for leaf 3 is given by eqn. (1) and dynamics of uninfected leaves is given by eqn. (3). Dynamics of infection in other leaves is described by the following equations:

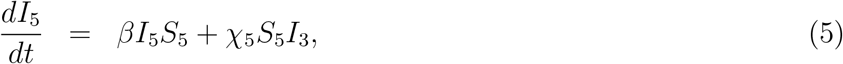

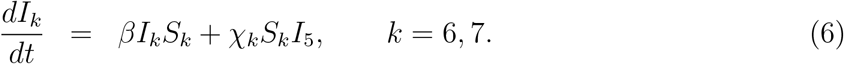 Initial conditions for the model are *I_k_*(0) = *I*_0_ if *k* = 3 and *I_k_*(0) = 0, otherwise, and *S_k_*(0) = 1.
- **Alternative Model 3.** Infection for leaf 3 is given by eqn. (1) and dynamics of uninfected leaves is given by eqn. (3). Leaf 3 is the only leave that contributes to infections of higher leaves. Dynamics of infection in other leaves is described by the following equations:

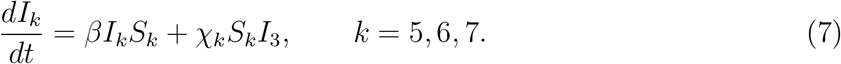 Initial conditions for the model are *I_k_*(0) = *I*_0_ if *k* = 3 and *I_k_*(0) = 0, otherwise, and *S_k_*(0) = 1.
- **Alternative Model 4.** The initial infection occurs on leaf 5 which contributes to infections of leaves 3, 6, and 7. All imported virions for these leaves comes exclusively from leaf 5. Dynamics of uninfected leaves is given by eqn. (3). Dynamics of infection in other leaves is described by the following equations (see Figure 1C):

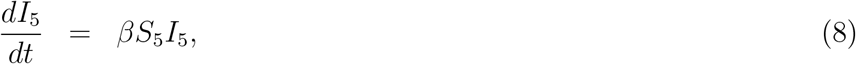

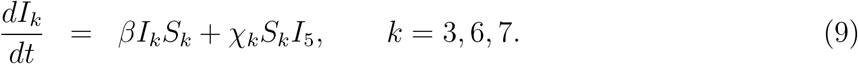 Initial conditions for the model are *I_k_*(0) = *I*_0_ if *k* = 5 and *I_k_*(0) = 0, otherwise, and *S_k_*(0) = 1.
- **Alternative Model 5.** The initial infection occurs on leaf 6 which contributes exclusively to the infections of leaves 3, 5, and 7. Dynamics of uninfected leaves is given by eqn. (3) and dynamics of infection in other leaves is described by the following equations:

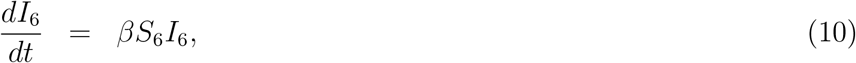

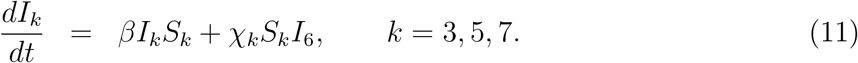 Initial conditions for the model are *I_k_*(0) = *I*_0_ if *k* = 6 and *I_k_*(0) = 0, otherwise, and *S_k_*(0) = 1.
- **Alternative Model 6.** The initial infection occurs on leaf 7 which contributes exclusively to the infections of leaves 3, 5, and 6. Dynamics of uninfected leaves is given by eqn. (3) and dynamics of infection in other leaves is described by the following equations:

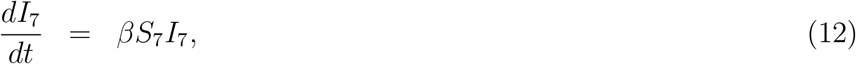

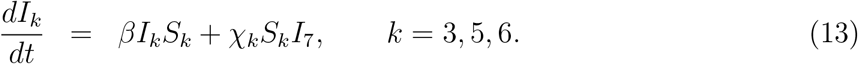 Initial conditions for the model are *I_k_*(0) = *I*_0_ if *k* = 7 and *I_k_*(0) = 0, otherwise, and *S_k_*(0) = 1.
- **Alternative Model 7.** The initial infection occurs on leaf 7 and virus accrues downward; it is essentially the model by Tromas *et al*. [1] being inverted. Dynamics of uninfected leaves is given by eqn. (3) and dynamics of infection in other leaves is described by the following equations (see Figure 1D):

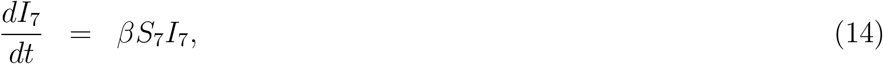

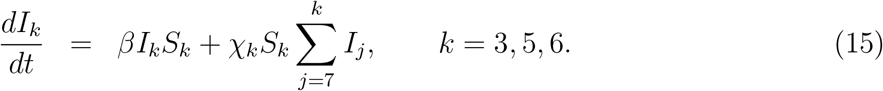 Initial conditions for the model are *I_k_*(0) = *I*_0_ if *k* = 7 and *I_k_*(0) = 0, otherwise, and *S_k_*(0) = 1.
- **Alternative Model 8.** The model assumes that infection starts in all leaves and proceeds independently (aka “logistic” model for individual leaves). Dynamics of infection in all leaves is described by the following equations:

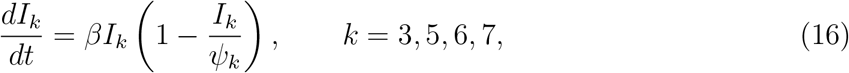 Initial conditions are *I_k_*(0) = *I*_0_*k*__, *k* = 3, 5,6, 7.
- **Alternative Model 9.** In all previous models virus dissemination within a given leaf stops when the fraction of infected cells reaches *ψ_k_* (e.g., eqn. (3)). This is also observed in the data. However, specific mechanisms of why the infection stops while not all cells in the leaf are infected were not fully investigated. Therefore, in our alternative model we assume that the dynamics of virus infection in a given leaf are not infection level-dependent but instead time-dependent. We define *T_k_* to be the time that the *k*^th^ leaf accumulates the “immune response” to stop the spread of the virus inside it, and *n_k_* represents how quickly this immune response kicks in. The dynamics of the infection is given by same equations as in the Tromas *et al*. [1] model (eqns. (1)–(2)), and the dynamics of uninfected cells available for infection due to generation of the immune response in the *k*^th^ leaf is given by

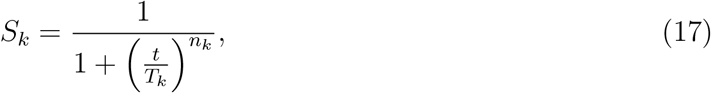

where the initial conditions for the model are *I_k_*(0) = *I*_0_ if *k* = 3 and *I_k_*(0) = 0, otherwise. This model has 4 extra parameters as compared to other alternative models but the model can be reduced in size by assuming that some of the parameters (e.g., *T_k_* or *n_k_*) to be leaf number-independent (see Main text for results).

#### Virus dissemination models for the infection/coinfection with two viral variants

In the experiments, the plants were infected with equal mixture of two viral variants, TEV-Venus and TEV-BFP [1]. However, the original model of Tromas *et al*. [1] and our previous alternative models did not discriminate between infection of the cells with two variants. The following alternative models now make this distinction. In these models we denote *V_k_* and *B_k_* as the fraction of Venus-infected and BFP-infected cells, respectively, and the fraction of coinfected cells is denoted as *M_k_*. Because our analysis illustrated that the specific pathway of TEV dissemination in 4 week old *N. tabacum* plants cannot be fully resolved using infection data alone we assume the dissemination pathway of Tromas *et al*. [1]. Then the dynamics of infection of plant leaves with the two viral variants we use the following equations:

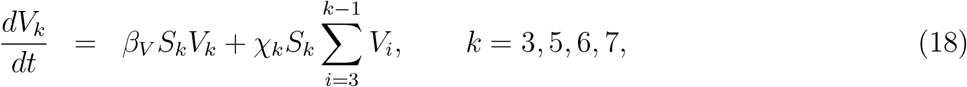

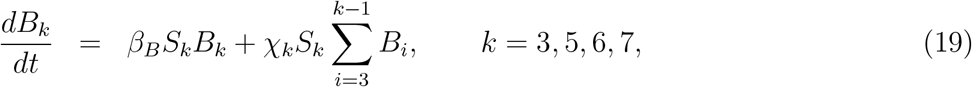

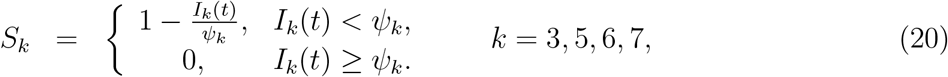

where *β_B_* and *β_V_* are the within-leaf infection rates fot BFP and Venus viruses, respectively, and *I_k_* (*t*) = *V_k_* + *B_k_* + *M_k_*. Note that we assume that virus dissemination to upper leaves is strain-independent. The initial conditions for all the following models are *V_k_*(0) = *V_0_, B_k_* = *B_0_*, and *M_k_*(0) = *M*_0_ if *k* = 3 and 0 otherwise. To describe the kinetics of viral coinfection we consider several alternative mathematical models.

- **1-alpha coinfection model.** In this model, we describe the coinfection growing as dependent on the within-leaf spread dynamics of both viruses. Here, and in other models *V_k_B_k_* is proportional to the rate at which coinfections are expected to arise by chance. We sum these these rates assuming that cells are first infected by one variant and then coinfected with another, and use a scaling factor *α* to indicate synergy (*α* > 1) or inhibition (*α* < 1) of the coinfection process as compared to random, mass action-like infection process:

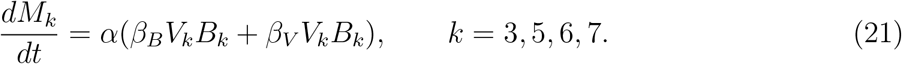
- **2-alpha coinfection model.** We assume that the rate of coinfection may proceed differently by the two viral strains denoted by *α*_1_ and *α*_2_ which is a simple extension of the 1-alpha coinfection model (eqn. (21)):

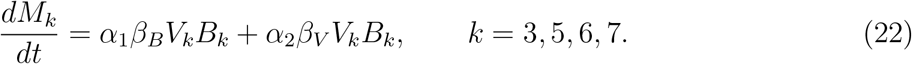
- **Probabilistic model.** Because *V_k_* and *B_k_* measure the fraction of cells infected by the particular virus in the *k*th leaf, then for fraction of coinfected cells, *M_k_*, we can think of the probability of a cell being infected by both strains as being determined by *B_k_ V_k_*. We can then use parameter *α* to measure how much more or less often coinfection is happening as compared with random chance: *α* =1 means coinfection is behaving like a random process; *α* < 1 means coinfection is occurring less often than it would by random chance, and *α* > 1 means coinfection is occurring with greater frequency than random chance [30]. Multiplying by *α* the product *B_k_V_k_* and differentiating it with respect to t gives:

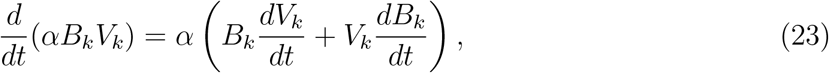

which then with the use of eqns. (18)–(19) results in the following model for the dynamics of coinfected cells

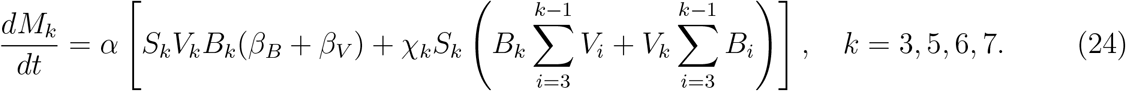 Note that in contrast with previous models (e.g., eqn. (22)), in this model coinfection within the leaf depends on the fractions of uninfected (*S_k_*) and virus-infected cells in the leaf (*V_k_* and *B_k_*).
- **2-alpha probabilistic model.** As in the original Tromas *et al*. [1] model, the equation for coinfection in the probabilistic model is composed of two parts (eqn. (24)): the first term with parameters *β_B_* and *β_V_* represents the within-leaf spread, and the second term with the parameter *χ_k_* represents the leaf-to-leaf spread. It seemed reasonable that coinfection may be driven more by one form of spread or the other, so we used *α*_1_ and *α*_2_ to measure their respective contributions:

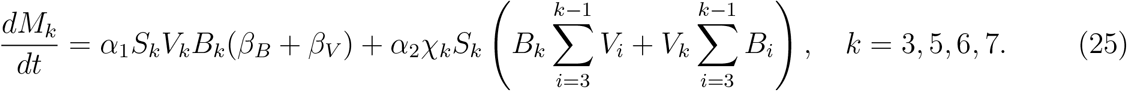
- **Logistic model for coinfection growth.** The details of how plant cells become coinfected by two different viruses during the local spread are not fully understood. Because typically plant viruses spread to adjacent cells via plasmadesmata, a coinfected cell may be source of both viral strains when infecting neighboring cells. In this alternative model we therefore assume that the frequency of coinfected cells increases randomly due to viral dissemination systemically from other leaves and logistically due to local, within-leaf spread:

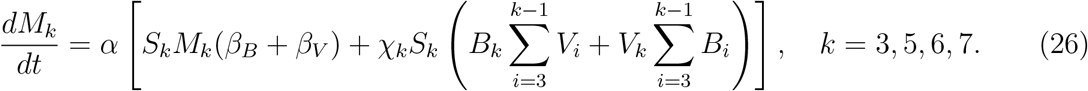
- **2-alpha logistic model for coinfection growth.** Similarly to the 2-alpha probabilistic model, the rate of coinfection may be different between local and systemic viral spread (eqn. (25)). Therefore, we use *α*_1_ and *α*_2_ to differentiate between coinfection occurring as within-leaf and leaf-to-leaf/systemic spread, respectively:

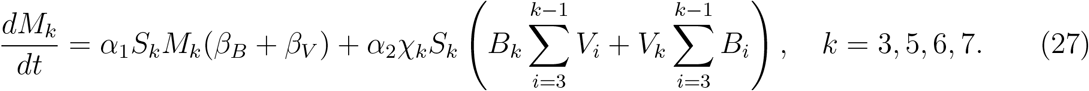

### Statistics

To fit models to data we used two alternative approaches. Tromas *et al*. [1] proposed to use the following binomial distribution-based likelihood to fit the models to data

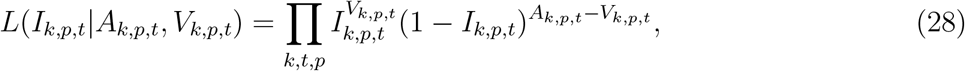

where *L* is the likelihood of the model given the data, *I_k,t,p_* is the model prediction for the frequency of infection (by either or both viral variants) of the particular leaf *k* and time point *t* of a plant *p, V_k,p,t_* is the number of infected cells observed in a sample, *A_k,p,t_* is the total number of cells observed in the sample (*k* is the leaf number, *p* = 1 … 5 is the plant replicate number, and *t* is the day on which the observation was made). The model parameters are estimated by minimizing the negative log likelihood (*nll*) given by the formula

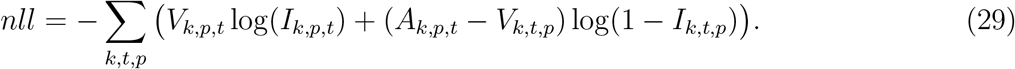

In the “coinfection” models we track the dynamics of cells infected with individual viral strains as well as coinfected cells. In these models *I_V_k,p,t__, I_B_k,p,t__, I_M_k,p,t__* represent the model predictions for the frequency of Venus- or BFP-infected, or coinfected cells, respectively. Therefore, to fit the coinfection models to data we extended the binomial distribution-based likelihood in the following way. We let *V_V_k,t,p__, V_B_k,t,p__, V_M_k,t,p__* be the number of cells infected by Venus, BFP, or both, respectively, as was measured experimentally. Note that *V_k,t,p_* = *V_V_k,t,p__* + *V_B_k,t,p__* + *V_M_k,t,p__*. Then we let

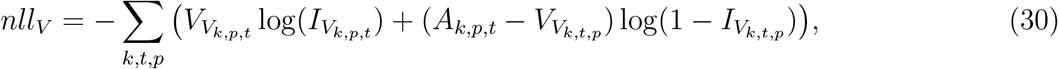

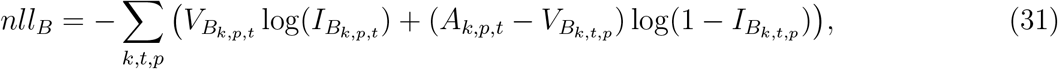

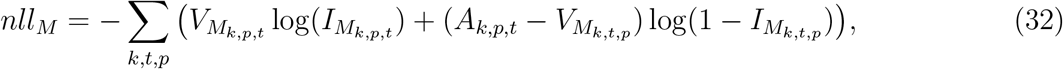

and nll is simply

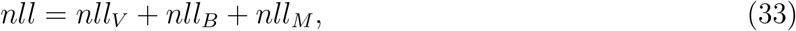

where the best fit parameters are found by minimizing the nll.

Binomial distribution-based likelihood takes into account the number of cells (protoplasts) extracted from each leaf. The total number of extracted cells varied dramatically between leaves (by up to 8 fold). It was therefore possible that different numbers of cells in the data may skew the likelihood-based estimates towards measurements with more cells. We therefore aimed to investigate whether other methods, e.g., assuming normally distributed data, i.e., normal distribution-based likelihood or least squares, can be used to fit the models to data. We tried several different ways of how least squares could be used to fit the model to data.

One approach is to use the frequency of infected cells *I_k,t,p_* as predicted by the mathematical model with the data *V_k,t,p_/A_k,t,p_*. For the models that only consider uninfected and infected cells (i.e., cells infected with either viral variant or coinfected with both variants), the sum of squared residuals (*SSR*) was then calculated as follows:

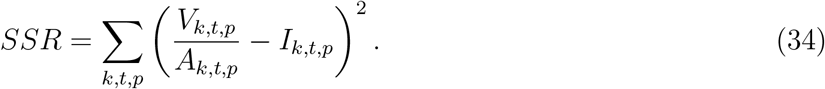

In our analyses we found that such a method does not typically result in normally distributed residuals (see Results section for details). Given large variability in the frequency of infected cells over time we applied log-transformation to the data and the model predictions and calculated the SSR using the following formula:

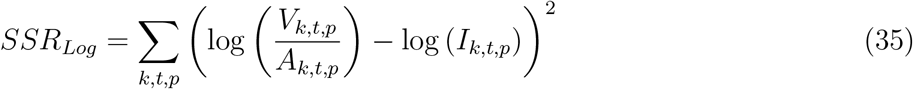

where the notations are the same as in eqn. (34). Log-transformation of the data, however, is problematic because in 2 cases of leaf 5 infection, the measured frequency of infected cells was 0. One approach was to remove such data points from the analysis but data removal can generate biases in the model fits, and therefore, we opted for a more appropriate approach whereby we replaced zeros in the data and the model predictions with the limit of detection (LOD). LOD in the data for infected cells was defined as the lowest value found in the data (for infected cells LOD = 5.12 × 10^−4^).

Similarly to eqn. (33) we used the following definition for SSR to fit the coinfection models to the data on the frequency of cellular infection with Venus (*V_V_k,t,p__*), BFP (*V_B_k,t,p__*) or both viruses (Mixed) (*V_M_k,t,p__*)

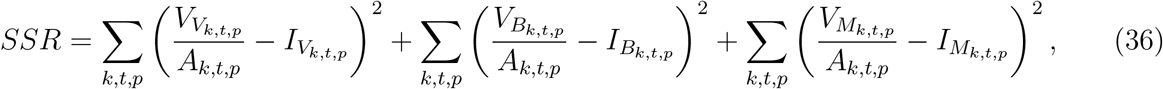

and the following is the below log-transformed variant (eqn. (37)):

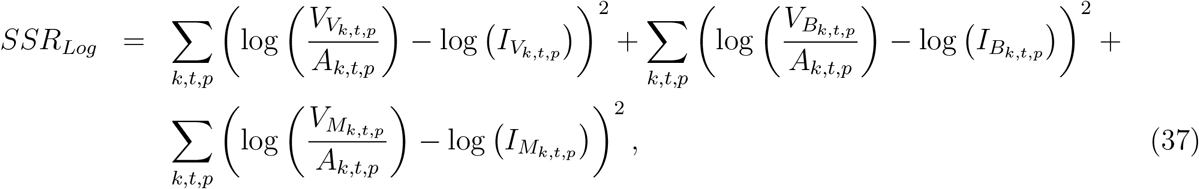

where data in which the frequency of infected cells was zero, we replaced these zero values with the LOD for individual virus-infected cells as LOD*_Venus_* = 8.43 × 10^−5^, LOD*_BFP_* = 3.26 × 10^−4^, and LOD*_Mixed_* = 3.2 × 10^−5^.

For binomial distribution-based likelihood, confidence intervals for best fit parameters were estimated by bootstrapping the data with replacement (sampling a given plant) 1000 times [31]. For least squares, we used routine `minimize` in from python library `lmfit` that provided 95% confidence intervals for estimated parameters.

To compare alternative mathematical models we used Akaike Information Criterion, *AIC*, that are calculated differently for binomial distribution- and normal distribution-based (least squares) likelihoods [28]:

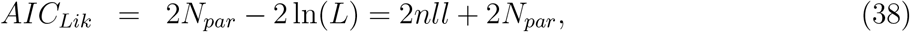

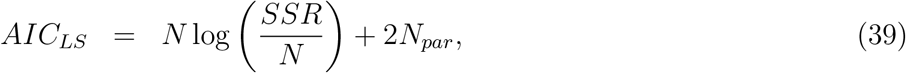

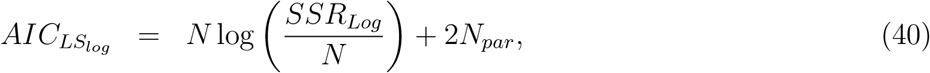

where *N* is the number of data points in the sample (in this case *N* = 80), and *N_par_* is the number of model parameters estimated by fitting the model to the data. Note that AIC differences of 0-4 are typically considered to be small while a difference of 10 indicates inferiority of the worse fit model in describing the data [28].

If plant cells are infected randomly by two different strains of the virus we expect that the frequency of coinfections with two viruses should be proportional to the product of the frequency of infections with single viral strains. To estimate the deviation from the random coinfection we used Odds Ratio of infection (*OR*) proposed previously to estimate deviation from random coinfection for HIV [30]:

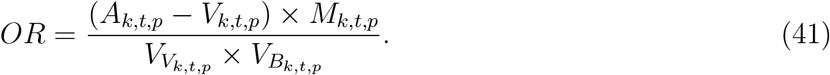

where *A_k,t,p_* – *V_k,t,p_* is the number of uninfected cells and *V_k,t,p_* = *V_V_k,t,p__* + *V_B_k,t,p__* + *V_M_k,t,p__* is the total number of infected cells in the data for the *k*^th^ leaf, time point *t*, and plant *p*.

### Programming details

All major analyses were done in Python (ver. 3.7.2) and some analyses were repeated in `R` (ver. 3.9.1). Python libraries used were `matplotlib` (ver. 3.3.2), `Pandas` (ver. 1.1.3), `NumPy` (ver. 1.19.0), `lmfit` (ver. 1.0.1), and `SciPy` (ver. 1.5.2). To solve the ODE-based models we used the odeint routine from `scipy.integrate` package. To fit models to data we used a differential evolution algorithm when the goodness of fit metric was *nll*, and when minimizing least squares residuals we used the Levenberg-Marquardt algorithm with a trust region. Both methods are part of Python’s lmfit library. To ensure reproducibility of our results as a part of this publication we share the data and the code to fit the original virus dissemination model to data using either binomial distribution-based likelihood or least squares, and the code to illustrate the impact of various parameters on the virus dynamics according to 2-alpha probabilistic model.

## Results

### Model with Tromas *et al*. [1] parameter values does not match the data

In their original study, Tromas *et al*. [1] manually introduced two different strains of TEV to the third leaf of the 4 week old *N. tabacum* plants and counted the number of infected and uninfected cells in different leaves (*k* = 3, 5, 6, 7) of the infected plants over time (*t* = 3, 5, 7, 10 days). Given that plant cells are immotile and each surrounded is by cellulosic cell walls, viruses can infect other cells in the leaf via two ways: 1) by passing through pores in the cells’ membranes and cell walls (called plasmodesmata) creating portals between adjacent cells, or 2) by entering the vasculature and migrating with phloem to other (sink) leaves of the plant [32]. Over time, the viral infection disseminates unequally between the leaves (Figure 3 and Figure S1). In particular, only about 10% of all cells in the originally inoculated leaf 3 become infected by 10 days of infection (Figure 3A), while on average 30% of cells become infected in leaves 6 and 7 (Figure 3C-D). Interestingly, leaf 5 becomes minimally infected (Figure 3B), and infection did not spread to leaf 4 [1]. There was great variability between infection of leaves in individual plants; for example, in leaf 7 by day 10 less than 10% of cells were infected in one plant while over 40% were infected in another plant (Figure 3D).

**Figure 3:**
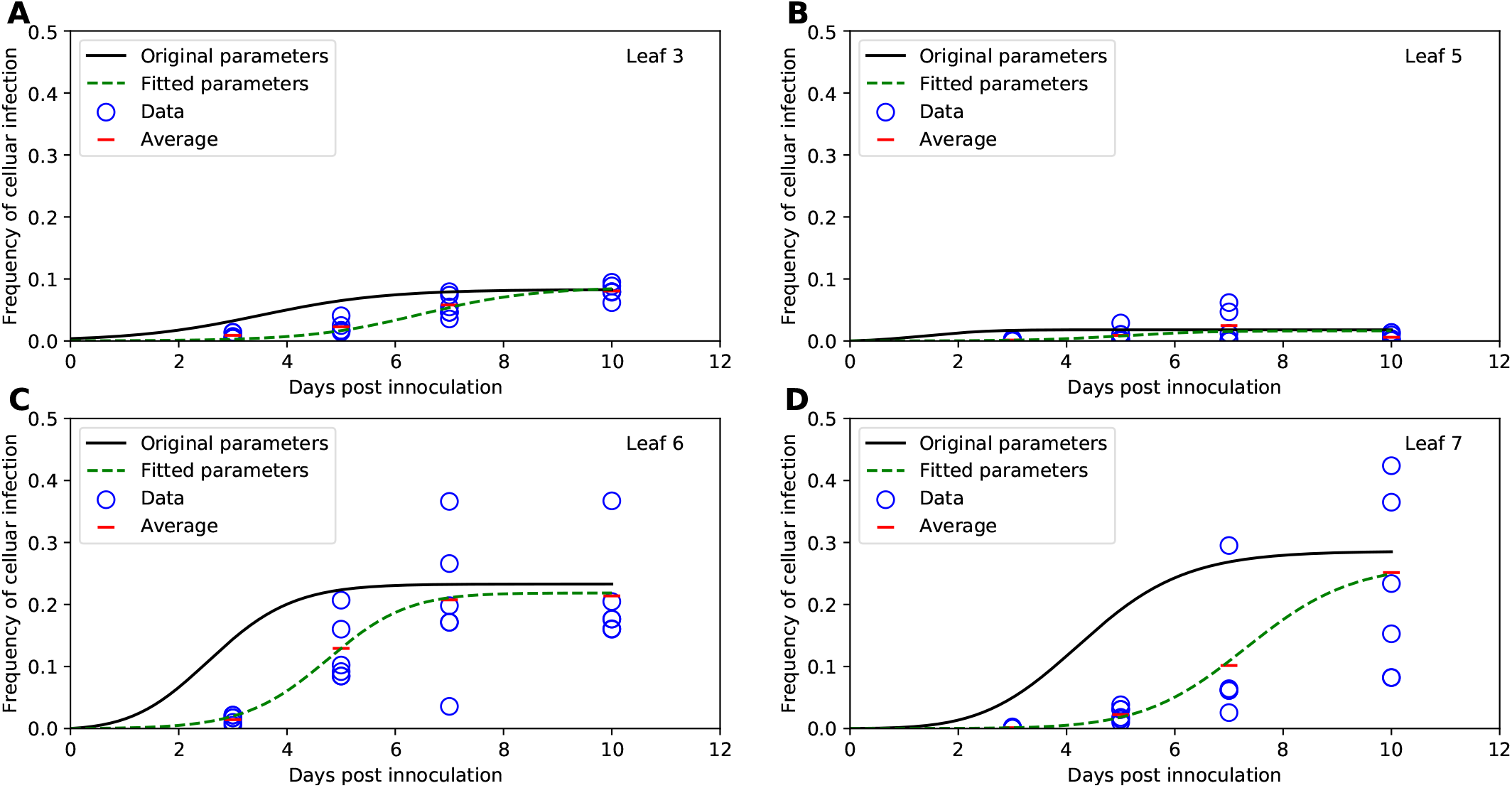
Previously published parameters estimates in Tromas *et al*. [1] do not provide a reasonable description of the data. We simulated basic mathematical model for viral spread in plants developed by Tromas *et al*. [1] (given in eqns. (1)–(3) and Figure 1A) using parameter values provided in the original publication (solid lines), or fitted the model to the data using binomial distribution-based likelihood method (eqn. (29), dashed lines). Data for the fraction of infected cells are shown by markers for leaf 3 (A), leaf 5 (B), leaf 6 (C), and leaf 7 (D) with red horizontal lines denoting average fraction of infected cells per time point. Parameters for the model fits are shown in Table 1.

To estimate basic parameters determining kinetics of TEV spread in *N. tabacum* plants, Tromas *et al*. [1] developed a mathematical model assuming that virus infection proceeds locally in each leaf and spreads from lower to upper leaves (Figure 1A). Via several model iterations, the model in which within-leaf virus spread was leaf number-independent but the virus transport to upper leaves from the lower leaves was leaf number-dependent, fitted the data with best quality [1].

To verify these results we simulated virus spread dynamics using Tromas *et al*. [1] published parameter values (Table 1) and the model equations (eqns. (1)–(3)) and compared model predictions with the data (provided by Tromas *et al*. [1]). Surprisingly, the model predictions did not match the average infection levels observed in the data (solid lines in Figure 3). While we did not fully know the exact reasons for this discrepancy, we found that if we were to shift the infection trajectories by 3 days, the model predictions matched the data relatively well (Figure S2). We therefore hypothesize that when numerically solving the model, Tromas *et al*. [1] may have initiated the solver starting at day 3 post infection given that it is the first time point at which experimental measurements were taken. (It is typical to obtain model predictions for times as given in the data, and solvers in R or python typically take the first time point as the time at which initial conditions are provided and not at the time 0 as is often assumed in models.)

**Table 1:** New parameter estimates for the basic mathematical model of virus spread in plants. We list the parameter estimates and 95% confidence intervals (CIs) of the basic mathematical model of viral spread in plants (eqns. (1)–(3)) as provided by Tromas *et al*. [1] (“Original parameters (*nll*)”) or by fitting the model to data in this work using using binomial distribution-based likelihood (“New parameters (nll)”, eqn. (29)) or using least squares with a logarithmic transform (“New parameters (*log LS*)”, eqn. (35)). Fits of the mathematical model for two sets of model parameters are given in Figures 3 and 4. Confidence intervals for best fit parameters were generated using bootstrap by resampling the data (for likelihood-based fits) or were provided by the routine `minimize` in from python library `lmfit` for least square-based fits (see Materials and methods for more detail).

|  | Original parameters (*nll*) |  | New parameters (*nll*) |  | New parameters (*log LS*) |  |
| --- | --- | --- | --- | --- | --- | --- |
| Parameter | Estimate | 95% CIs | Estimate | 95% CIs | Estimate | 95% CIs |
| $I_0$ | 0.00372 | (0.001,0.017) | 0.00022 | (0.00011,0.00108) | 0.0005 | (0.0001,0.0008) |
| $\beta$, 1/day | 0.871 | (0.257,1.66) | 0.950 | (0.549,1.183) | 0.902 | (0.730,1.618) |
| $\chi_5$, 1/day | 0.724 | (0.033,0.813) | 0.167 | (0.042,17.291) | 0.063 | (0.023,0.162) |
| $\chi_6$, 1/day | 1.38 | (0.580,2.340) | 1.046 | (0.228,1.487) | 0.691 | (0.387,1.053) |
| $\chi_7$, 1/day | 0.107 | (0.050,0.263) | 0.029 | (0.009,0.160) | 0.029 | (0.015,0.051) |
| $\psi_3$ | 0.083 | (0.053,0.147) | 0.080 | (0.074,0.134) | 0.074 | (0.051,0.096) |
| $\psi_5$ | 0.018 | (0.002,0.050) | 0.016 | (0.005,0.024) | 0.006 | (0.004,0.010) |
| $\psi_6$ | 0.233 | (0.155,0.345) | 0.224 | (0.203,0.287) | 0.204 | (0.181,0.234) |
| $\psi_7$ | 0.286 | (0.234,0.346) | 0.269 | (0.130,0.418) | 0.224 | (0.092,0.557) |

To check that the virus dissemination model of Tromas *et al*. [1] is consistent with experimental data we fitted the model to the data using binomial distribution-based likelihood (see Materials and Methods for more detail). Importantly, the model fitted the data visually with good quality (dashed lines in Figure 3) indicating consistency of the model with the data. Interestingly, while some model parameters, such as *ψ_k_* were similar in the original and corrected analysis such, others such as *I*_0_ or *χ_k_* differed substantially (Table 1). While confidence intervals for newly estimated parameters of the Tromas *et al*. [1] model are a bit large, we found that there is large difference in *AIC_Lik_* for this model when used with previously published Tromas *et al*. [1] parameters and our new estimates (Δ*_Lik_* > 100, results not shown). Thus, our analysis provided updated and correct estimates of parameters characterizing kinetics of TEV spread in *N. tabacum* plants in the Tromas *et al*. [1] model.

### Fitting the models using binomial distribution-based likelihood or normal distribution-based likelihood (least squares) delivers similar parameter estimates

In their study, Tromas *et al*. [1] proposed to use binomial distribution-based likelihood to fit the models to data. In this approach, the probability of a plant cell to be infected was treated as a Bernoulli trial in which *A* total cells are sampled, and the number of infected cells (*V*) is determined. While it seemed reasonable it was not fully justified why such a likelihood is a good choice. There may be several potential issues with it. First, because there was a large variability in total number of cells recovered from different leaves (from minimal 4314 to maximal 32168 protoplasts/leaf), the data are unbalanced. Sources of such variability, however, are not entirely clear and may be due to different sizes of different leaves but also may be related to difficulty of isolating protoplasts from leaves [33]. Parameter estimates may be biased if the fit favors better description of the data with the larger number of isolated cells. Second, while the large number of cells isolated may indicate certainty in estimation of the frequency of infected cells in a sample, there is a great variability in frequency of infected cells in the same leaf number between individual plants (e.g., Figure 3D), and binomial distribution-based likelihood may not take such variability adequately into account. Third and finally, given that a relatively large number of cells was measured in each leaf (> 10^3^), the distribution of the fraction of infected cells per central limit theorem may approach normal distribution, and therefore, one could use a normal distribution-based likelihood (least squares) for fitting models to data.

Therefore, we fitted the Tromas *et al*. [1] model (eqns. (1)–(3)) to the data using several different versions of least squares (see eqns. (34)–(35) and Materials and methods for more detail). Surprisingly, independently of the method used, the model predictions of the binomial distribution-based fits or least squares fits were nearly identical (e.g., Figure 4) and with a minimal, statistically nonsignificant difference in the parameter estimates for both fits (Table 1). Therefore, this result suggests that it may be reasonable to use least squares (or more generally, normal distribution-based likelihood) to fit virus dissemination models to these data. We did, however, find that not all least squares-based methods were appropriate. In particular, least squares with the frequency of infected cells resulted in skewed, non-normally distributed residuals (Shapiro-Wilk test, *W* = .785, *p* = 1.887× 10^−9^). Some of the traditional approaches, for example the arcsin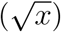 transformation for the frequency of infected cells did not normalize the residuals (*W* = 0.803, *p* = 5.745 × 10^−9^), however, log-transformation in which zero values were replaced with the limit of detection (LOD, see Materials and methods for more detail) did (*W* = 0.963, *p* = 0.021). Therefore, this analysis suggests that log-transformation of the data and model predictions is a viable alternative to the binomial distribution-based likelihood method of Tromas *et al*. [1].

**Figure 4:**
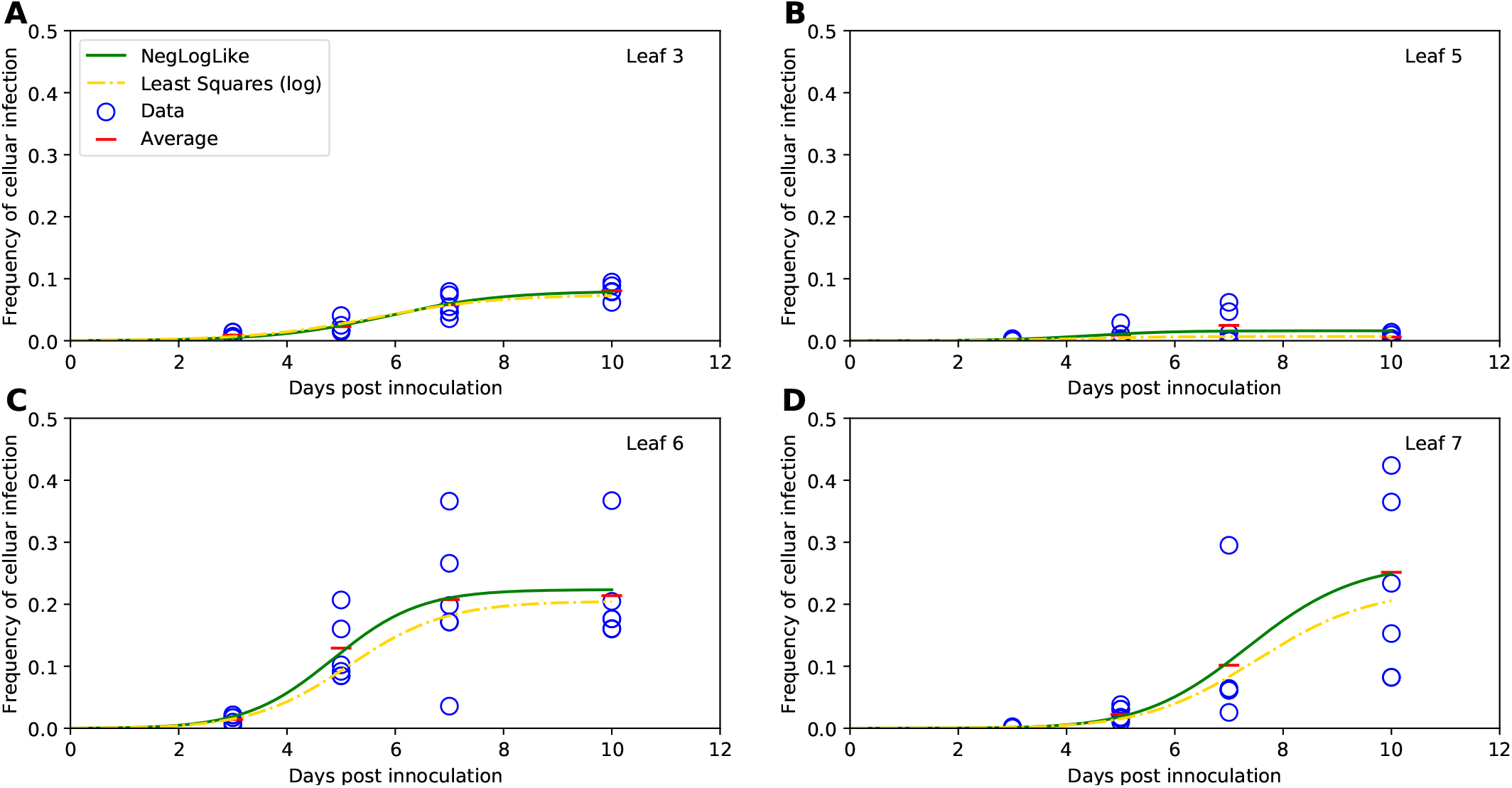
Best fits of the basic mathematical model found using either binomial distribution-based likelihood or least squares are nearly identical. We used either binomial distribution-based likelihood method (eqn. (29), [1]) or least squares for log-transformation of the data and model predictions (eqn. (35)) to fit the basic mathematical model (eqns. (1)–(3)) to the virus spread data for leaf 3 (A), leaf 5 (B), leaf 6 (C) and leaf 7 (D). Data on proportion of virus-infected cells are shown by markers and lines are the predictions of best fit models. Parameters for the model fit using likelihood and least squares with the log transform are given in Table 1 (“New parameters (*nll*)” and “New parameters (*Log LS*)” columns, respectively). In fitting the models using least squares for log-transformed data, the limit of detection (LOD) was LOD = 5.12 × 10^−4^.

### Several alternative models are consistent with viral spread kinetics

In their analysis Tromas *et al*. [1] investigated which parameters of the virus dissemination model may vary with the leaf number and by comparing alternative models they found that *ψ* and *χ* must be leaf-dependent to explain the data accurately. However, in that analysis they did not investigate if difference in virus dissemination may be due to variable within-leaf replication kinetics, determined by the parameter *β*, and not due to virus dissemination rate *χ*. Interestingly, we found that the alternative model (based on eqns. (1)–(3)) in which *β_k_* and *ψ_k_* vary with the leaf number *k* (i.e., virus dynamics in a given leaf is determined mainly by the local spread in the leaf) while systemic dissemination of the virus to upper leaves is constant (*χ_k_* = *χ*) fitted the data with similar quality (as judged by *SSR* or *AIC*) as the original model. This alternative model has an extra parameter because of four *β* for four leaves studied while in the original model *χ* was defined for three leaves only. Yet, this result already suggested that the data on variable virus accumulation in different leaves can be explained equally well by differences in how much virus is delivered to upper leaves (*χ_k_*) or by differences in how the virus replicates and spreads in individual leaves (*β_k_*).

We next questioned whether a specific pattern of virus dissemination from the inoculated leaf 3 to the upper leaves can be determined from these experimental data. While there is a general understanding of how viruses in plants disseminate after a local infection (e.g., [25]) details of the dissemination may vary by the plant species, age, conditions in which the plant was grown, the virus species, inoculation method, and many other details. For example, the time when individual leaves become sources or sinks for sugar transport – which will influence virus dissemination pathways – depends on many environmental and developmental factors [26]. Because many of these details are unknown for a specific experimental set-up we investigated if the information provided by experimental data on the fraction of infected cells in individual leaves over time is sufficient to establish a pattern for systemic viral dissemination.

Therefore, we developed a series of alternative mathematical models in which the pattern of virus dissemination differed in multiple ways from the original dissemination model of Tromas *et al*. [1] (Figure 1B-D and eqns. (4)–(17)) and fitted these models to data. For example, alternative model 1 assumed that virus dissemination to upper leaves occurs only from the leaf below it, i.e., from leaf 3 to leaf 5, and then from leaf 5 to leaf 6 and so on (Figure 1B). Alternative model 7 assumed that even though virus inoculation occurred at leaf 3, somehow virus immediately disseminated to leaf 7, and then spread to lower leaves (Figure 1D and see Materials and Methods for details for other models). These alternative models should not be necessarily considered as inappropriate, because, for example, at day 3 after infection, on average leaf 6 had already nearly twice as many infected cells as the leaf 3 (0.014 vs. 0.009).

Finding the best fit model depended strongly on the statistical method used for fitting models to data. For example, using binomial distribution-based likelihood method suggested that best fit is provided by the alternative model 2 with the Tromas *et al*. [1] model fitting the data significantly worse (Table S1, Δ = 290). We hypothesize that this result arose because of high sensitivity of such likelihood function to the experimental measurements, especially at the low frequency of infected cells. In contrast, fitting the models to data using least squares (eqn. (34)) provided fits of all models with identical quality (results not shown). This result was driven by the need of the models to more accurately fit the data with high frequency of infected cells in leaves 6 and 7 at later time points, at the expense of poorer fits of other data. These fits, however, were not adequate due to non-normally distributed residuals as was observed when fitting Tromas *et al*. [1] model to data (see above). Finally, fitting the models to log-transformed data (and replacing the zero values with the LOD) provided a more graded classification of alternative models (Table 2). In particular, three models (original and alternative models 1&2) assuming that virus dissemination starts from leaf 3 provided better fits (based on AIC) than the models assuming that spread starts from upper leaves (e.g., alternative model 7). Interestingly, the quality of the model fits deteriorated as the models assumed virus dissemination not to originate from leaf 3 — i.e., the models in which dissemination started at leaf 5 or 6 fitted the data with better quality than the model in which dissemination started at leaf 7 (Table 2). This result suggests that the data on virus dissemination does contain the signal indicating the virus most likely starts spreading from leaf 3 upwards; however, the strength of such statement from our mathematical modeling-based analysis is relatively weak. Thus, these experimental data do not provide strong evidence for a specific route of TEV dissemination in *N. tabacum* plants. There are some good news, however. Some parameters appear to be robustly estimated in all the models such as *β* and *ψ_k_*; that is perhaps unsurprising given that these parameters determine within-leaf viral spread.

**Table 2:** Several alternative models provide somewhat different fits of the virus spread data with different parameter sets. We fit seven alternative models for virus spread kinetics (given in eqns. (4)–(15)) to the data on viral spread in plants using least squares with a logarithmic transform (see eqn. (35)). Along with parameter values for every model we provide the total error (*SSR_Log_*), *AIC_LS_Log__*, and Δ*_AIC_* (difference in AIC between the model with the lowest AIC and all other models). We also show the results of the Shapiro-Wilk normality test (*W* and *p* value) applied to the residuals of the fitted models.

| Fitted with least squares method (Log transformation & 0 ≡ LOD) | | | | | | | | |
| --- | --- | --- | --- | --- | --- | --- | --- | --- |
| Parameter | Original | Alt.<br>Model 1 | Alt.<br>Model 2 | Alt.<br>Model 3 | Alt.<br>Model 4 | Alt.<br>Model 5 | Alt.<br>Model 6 | Alt.<br>Model 7 |
| $I_0$ | 0.0005 | 0.0008 | 0.0006 | 0.0001 | 0.00006 | 0.0008 | 0.0005 | 0.0005 |
| $\beta$, 1/day | 0.902 | 0.744 | 0.871 | 1.299 | 1.050 | 1.028 | 1.116 | 1.159 |
| $\chi_3$, 1/day | N/A | N/A | N/A | N/A | 1.672 | 0.133 | 1.541 | 0.286 |
| $\chi_5$, 1/day | 0.063 | 0.059 | 0.059 | 0.322 | N/A | 0.025 | 0.277 | 0.050 |
| $\chi_6$, 1/day | 0.691 | 8.201 | 8.025 | 3.987 | 3.663 | N/A | 3.079 | 3.263 |
| $\chi_7$, 1/day | 0.029 | 0.073 | 0.749 | 0.477 | 0.332 | 0.026 | N/A | N/A |
| $\psi_3$ | 0.074 | 0.083 | 0.075 | 0.072 | 0.060 | 0.059 | 0.055 | 0.051 |
| $\psi_5$ | 0.006 | 0.006 | 0.006 | 0.005 | 0.006 | 0.006 | 0.005 | 0.005 |
| $\psi_6$ | 0.204 | 0.204 | 0.199 | 0.194 | 0.201 | 0.215 | 0.189 | 0.184 |
| $\psi_7$ | 0.224 | 0.228 | 0.238 | 13.000 | 0.215 | 0.211 | 0.218 | 0.209 |
| $SSR_{Log}$ | 52.713 | 51.991 | 52.255 | 55.612 | 54.468 | 55.148 | 55.486 | 56.309 |
| $AIC_{SSR_{Log}}$ | -15 | -16 | -16 | -11 | -13 | -12 | -11 | -10 |
| $\Delta_{AIC}$ | 1 | 0 | 0 | 5 | 3 | 4 | 5 | 6 |
| $W$ | 0.963 | 0.959 | 0.960 | 0.973 | 0.969 | 0.973 | 0.972 | 0.973 |
| $p$ | 0.021 | 0.012 | 0.013 | 0.099 | 0.049 | 0.091 | 0.080 | 0.084 |

We tested two additional alternative models for how well they may describe these data. Alternative model 8 assumed that upon virus inoculation, virus disseminates to all leaves and then replicates in individual leaves independently of other leaves, as describe by logistic equation (eqn. (16)). Alternative model 9 assumed that reduction in the fraction of susceptible cells in a leaf is not determined by the fraction of infected cells but by the time since infection (eqn. (17)). The rationale for this modification is that it is possible that infection induces generation of a local or systemic immune response after a delay *T_k_* which renders uninfected cells resistant to infection. Both of these alternative models fitted the data well based on *SSR* or AIC metrics (Figure S3 and Table S2). Interestingly, the time-dependent cell susceptibility model suggested that differences in how quickly cells become resistant is leaf-dependent (Table S2); however, this could be due to differences in time of when immune response is triggered and/or “arrives” to a given leaf (determined by *T_k_*) or the speed at which uninfected cells in the leaf are rendered resistant (determined by *n_k_*). Taken together, our results strongly suggest that multiple pathways of TEV dissemination and growth in individual leaves in the *N. tabacum* plants are consistent with the data and additional experiments and/or data need to be involved to eliminate unreasonable models [34].

### Modeling dynamics of coinfection by two TEV strains

#### Odds ratio test implies a higher than random rate of coinfection

Our modeling-based analysis so far and that of Tromas *et al*. [1] treated cells in our data as infected or uninfected. However, in their experiments Tromas *et al*. [1] measured the fraction of cells infected with either or both of two viral strains of TEV, Venus or BFP (see Materials and Methods for more detail). Virus coinfection may impact many facets of viral dynamics and growth. A paramount consequence of two or more virions infecting the same cell simultaneously is that it may result in production of recombined variants as has been documented for human immundodeficiency virus (HIV) [35, 36]. In particular, in acute HIV infection, variants representing recombinants of infecting/founding strains, arose rapidly within a few months [37]; interestingly, a simple mathematical model predicted that accumulation of the variants can be simply due to random coinfection of the susceptible cells by two viral variants. Dang *et al*. [30] investigated whether infection of CD4 T cells in culture *in vitro* occurs randomly by two different HIV variants, HIV-eGFP and and HIV-IHSA. The authors proposed an odds ratio (*OR*) metric to estimate deviation of the rate of cell coinfection with two viruses as compared to single infections (eqn. (41)). Interestingly, in all theirs experiments with 2 HIV strains and different types of target T cells *OR* > 1 (typically, *OR* = 2 – 8), suggesting that coinfections were observed more often than single infections [30]. The authors explained this result by variability in CD4 T cell susceptibility to infection with susceptible cells being more easily infected with the two variants. A similar result was found later in another study [38]. Given our rich dataset on the dynamics of coinfection of plant cells with two variants of TEV we calculated the *OR* (eqn. (41)) for every leaf and every time point in our data.

Interestingly, we found very high values for *OR* for most of the data, all exceeding one, with many values being in the range 10-100 (Figure 5). Note that in some cases, mostly for leaf 5, we could not calculate *OR* due to absence of coinfected cells (Figure 5B). OR of 10 to 100 is much higher than that found previously for HIV [30]. There may be several reasons for that. First, it is possible that there is high degree of variability in susceptibility of different plant cells to infection, and cells that are highly susceptible get infected with both variants easily. We also found that there is a significant decline in *OR* with time of infection for all but leaf 5; this decline is consistent with the hypothesis that initially highly susceptible cells are infected resulting in high *OR* which declines as more resistant cells are infected (Figure 5). Alternatively, the mode of virus transmission within the leaf played the major role. Indeed, in plants viruses are transmitted from the infected cell to adjacent cells via plasmodesmata, and if a cell is coinfected with two variants, it is possible that all new infections occur by both viruses simultaneously [32]. Finally, if infection of cells occurs sequentially, infection with one variant may suppress any potential antiviral activity in the cell, allowing that cell to be coinfected with another variant [39–41]. To further test these hypotheses we used mathematical modeling.

**Figure 5:**
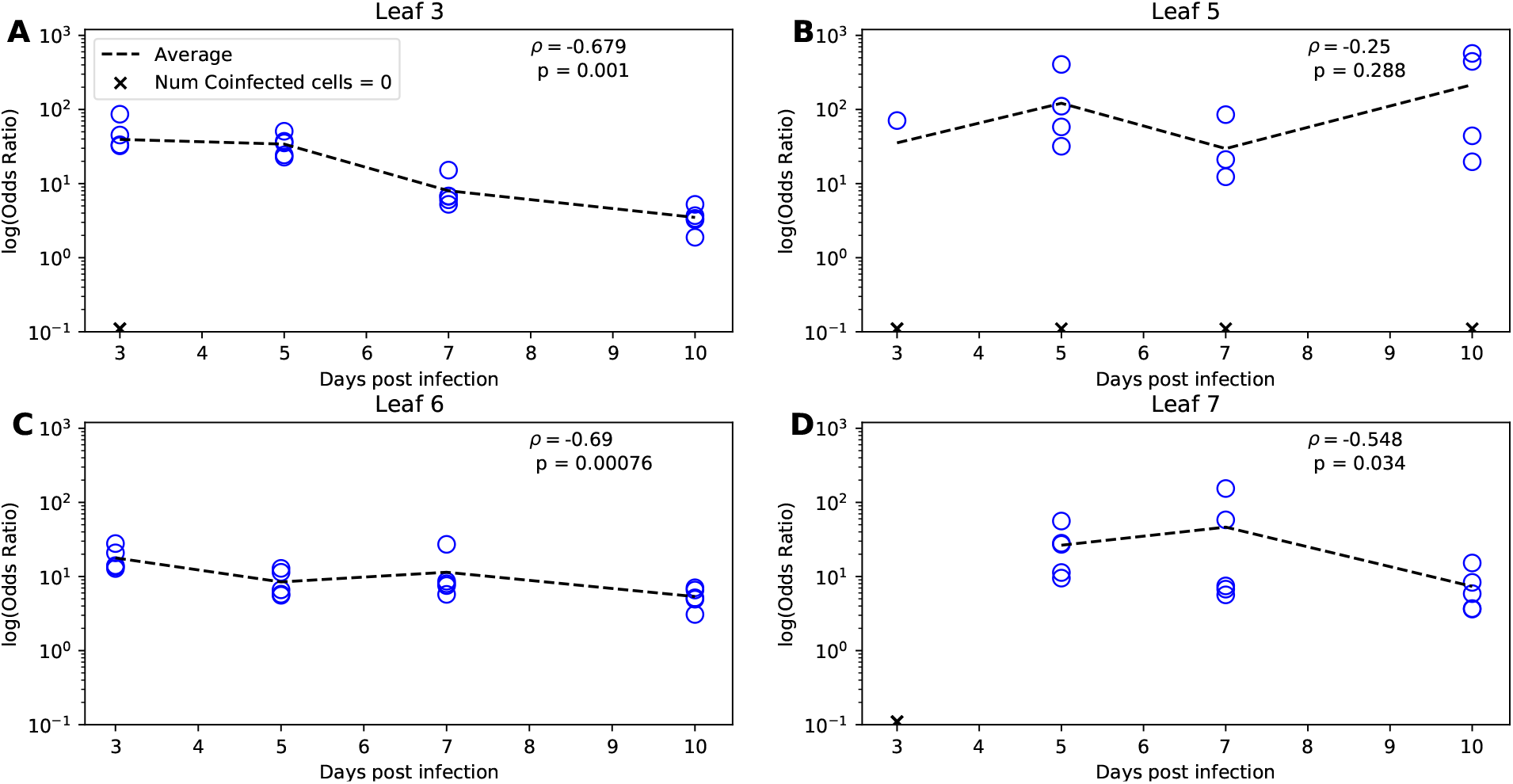
There is a high degree of coinfection of individual leaf cells by two different viruses. For each leaf we calculated the odds ratio (the relative probability of a cell being coinfected by two different viruses as compared to infection rate of cells by individual viruses, eqn. (41)) using a previously published method [30]. Note that when infection proceeds randomly, expected odds ratio is 1. Resulting odds ratio for individual plants are shown for leaf 3 (A), leaf 5 (B), leaf 6 (C), and leaf 7 (D); lines connect the average values per time point. Missing values (when odds ratio could not be calculated) are denoted as crosses. Spearman-Rank correlation *ρ* of the change in odds ratio with time per leaf and p-values from the test (*ρ* = 0) are shown on individual panels (when calculating *ρ* missing values were excluded).

#### Probability-based coinfection model performs best compared to other coinfection models

Given that many alternative mathematical models are consistent with the pathway of systemic virus dissemination (Table 2) to investigate potential mechanisms of TEV coinfection dynamics in different leaves we decided to fix the details of virus dissemination between leaves to those provided in the previous study [1], i.e., we let the virus infection to be initiated in the leaf 3 and dissemination to upper leaves to depend on the infection rate of leaves below (Figure 1A and eqns. (1)–(3)). To describe how coinfected cells are generated we developed six alternative “coinfection” models (see Figure 2 for 2 examples, eqns. (21)–(27), and Materials and Methods for more detail). In the first, 1-alpha coinfection model, dynamics of coinfected cells are driven only by the within-leaf frequency of cells infected with either of two variants with the parameter *α* determining deviations of the coinfection from random (Figure 2A and eqn. (21)). A simple extension of this model was to allow for different efficacies of coinfection depending of which virus infected the susceptible cell first (eqn. (22)). Two other models assumed that coinfection may happen via two different pathways: local, within-leaf infection dependent on the frequency of single-infected cells and uninfected cells and via between leaf virus dissemination, with either identical (*α*) or different (*α*_1_ and *α*_2_) weights for this coinfection processes (Figure 2B and eqns. (24)–(25)). Finally, the third set of two models assume that coinfection due to within-leaf dynamics occurs due to coinfected cells transmitting both viral variants to susceptible cells, and due to between-leaf dynamics occurs similarly as in the previous model. We similarly assume that these two processes may proceed with different deviations from random process which is captured by parameters *α*_1_ and *α*_2_ (eqns. (26)–(27)).

We fitted these models to experimental data using two alternative approaches, log-transformed least squares (with LOD replacements of zero values) and binomial distribution-based likelihood, both extended to account for singly and co-infected cells in each leaf (see eqn. (37) and eqn. (33) in Materials and methods for more detail). The 2-alpha Probabilistic model (eqn. (22)) was the best performing model when fitted by either method (Table 3). Importantly, with both methods the basic models assuming that coinfections occur randomly, due to within-leaf coinfection of cells poorly described the data (Table 3 and Figure S4).

**Table 3:** The 2-alpha probabilistic model fits the coinfection data with best quality. We fitted a series of mathematical models (see Materials and methods and Figure 2) that make different assumptions on how coinfection of individual cells with two different viruses occur to the data on viral spread. The models were fitted using the binomial distribution-based likelihood method (eqn. (33)) or the least squares method with a log transform of the data (eqn. (37)). AICs were calculated using eqn. (38) and eqn. (40), for the likelihood and least squares methods respectively. Values for nll, *AIC_Lik_*, and *AIC_LS_Log__* rounded to the nearest whole number. Δ*_AIC_* for both methods are calculated by taking the AIC score from the model and method in question and subtracting it from the lowest AIC in its corresponding row. In fitting models using least squares to log-transformed data we used the following values for the limit of detection: LOD*_Venus_* = 8.43 × 10^−5^, LOD*_BFP_* = 3.26 × 10^−4^, and LOD*_Mixed_* = 3.2 × 10^−5^.

| Parameters | 1-a Prob.<br>eqn. (24) | 2-a Prob.<br>eqn. (25) | 1-a Coin.<br>eqn. (21) | 2-a Coin.<br>eqn. (22) | 1-a Log.<br>eqn. (26) | 2-a Log.<br>eqn. (27) |
| --- | --- | --- | --- | --- | --- | --- |
| $nll$ | 512112 | 511728 | 519132 | 525576 | 515892 | 517656 |
| $AIC_{Lik}$ | 1024246 | 1023480 | 1038286 | 1051176 | 1031806 | 1035336 |
| $\Delta_{AIC}$ | 766 | 0 | 14806 | 27696 | 8326 | 11856 |
| $SSR_{Log}$ | 339.459 | 269.521 | 335.169 | 514.719 | 421.662 | 477.806 |
| $AIC_{SSR_{Log}}$ | 109 | 56 | 106 | 211 | 161 | 193 |
| $\Delta_{AIC}$ | 53 | 0 | 50 | 155 | 105 | 137 |

We also fitted the models using the least squares method for raw, untransformed frequencies of infected cells, but these fits poorly described the dynamics of coinfected cells (results not shown). We reasoned that this is because there are typically fewer coinfected cells than single-infected cells, and this least squares method favored fitting the dynamics of single-infected cells with better quality (due to their higher abundance). In this specific case, a statistical model based on untransformed least squares does not appear to be adequate.

With both of the appropriate methods we found that the 2-alpha probabilistic model fits the data with best quality, and the next best, 1 alpha probabilistic model performed significantly worse (per AIC scores, Table 3). Indeed, the model could very accurately describe the dynamics of single- and co-infected cells and predicted a more rapid increase in the coinfected cells for leaf 6 and 7 than that for single-infected cells (Figure 6). Unfortunately, we found relatively wide confidence intervals for estimates of many of these parameters except *ψ_k_* suggesting that the amount of data available was relatively low, and increasing the number of time points and/or plant repeats may have allowed for more precise estimates. We should note, however, that mean estimates for within-leaf infection rates *β_V_* and *β_B_* and between-leaf spread rates *χ_k_* were very similar to those found when fitting Tromas *et al*. [1] model to the data on infected cell dynamics (Table 1) lending some support that our model parameters are not unrealistic. Excitingly, we found that for within-leaf virus spread, coinfection rate was much higher than cell infection by single viruses (*α*_1_ = 10.1) supporting our analysis using odds ratio (Figure 5). The between-leaf coinfection rate was not different from random model (*α* ≈ 1) suggesting that most of coinfections were driven by within-leaf dynamics and not due to transfer of viruses systemically. This, perhaps, makes sense because locally it is easier for one cell to be coinfected by 2 viruses while when viruses enter the leaf at random locations due to systemic dissemination, coinfection is expected to be rare.

**Figure 6:**
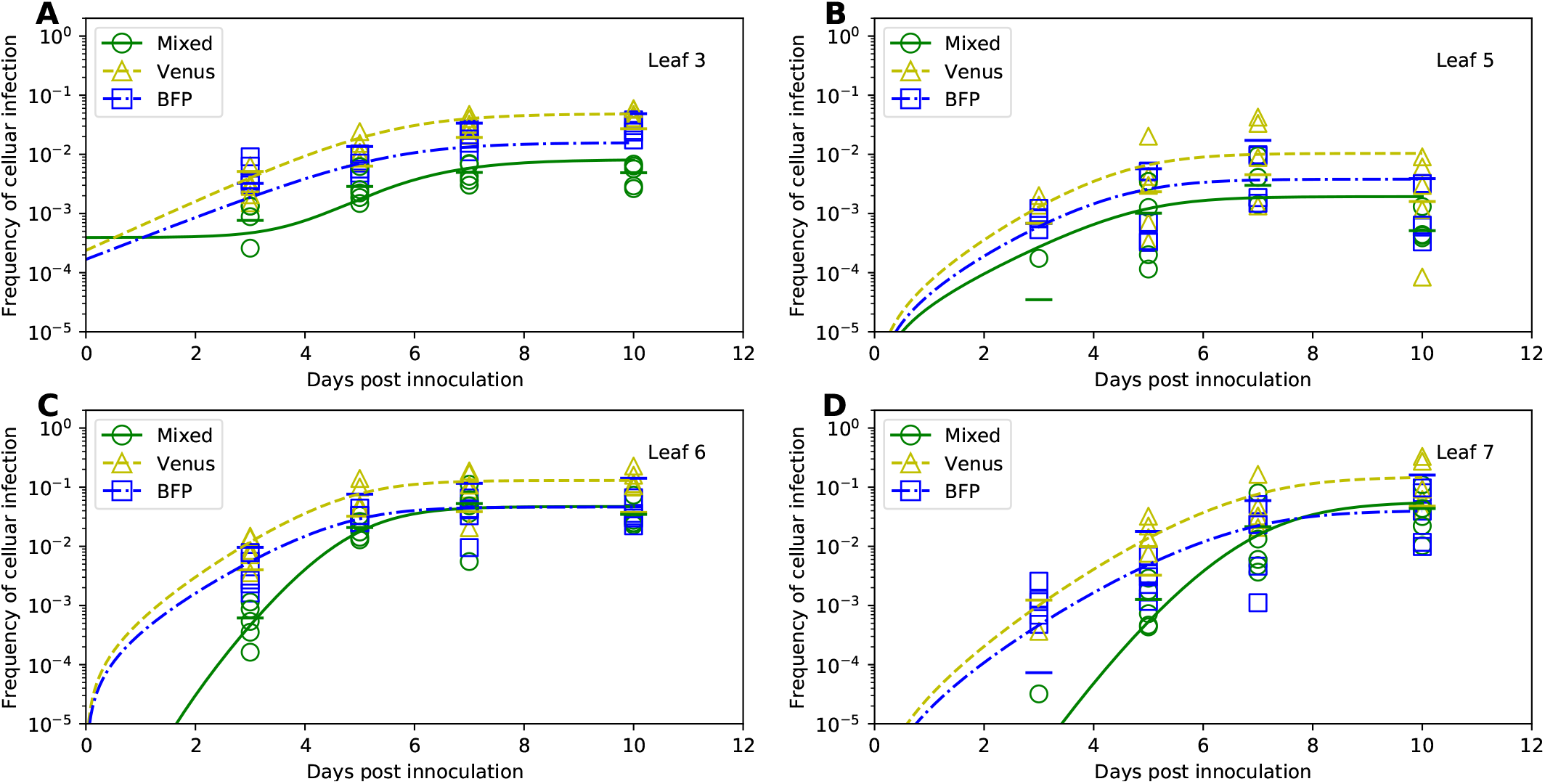
The 2-alpha Probabilistic Model fits the coinfection data with best quality. The 2-alpha Probabilistic Model (eqns. (18)–(19) and eqn. (25)) assumes that coinfection of individual cells by two different strains depends on the level of uninfected cells in the leaf (*S_k_*) and that local coinfections in the leaf occur at different kinetics that coinfection between leaves (Figure 2B). We fitted this model to the data on infection of cells by either individual viruses or coinfection of the same cell by different viruses. The model was fitted using the binomial distribution-based likelihood method (eqn. (29)). Markers show frequency of cells infected with Venus or BFP viruses or coinfected with both viruses (“Mixed”), and lines are predictions of the mathematical model. The short horizontal bars show the average infection rate for a given virus variant for a particular day and infected cell type. The parameters providing the best fit and their 95% confidence intervals (estimated using by boostrapping the data) are as follows: *V*_0_ = 0.0002 (2 · 10^−5^,0.001), *B*_0_ = 0.0002 (2 · 10^−8^, 0.001), *M*_0_ = 0.0008 (0.0, 0.001), *β_V_* = 0.975 (0.606,10)/day, *β*_B_ = 0.835 (0.426,10)/day, *χ*_5_ = 0.116 (0.004,9.522)/day, *χ*_6_ = 0.858 (0.0001,10)/day, *χ*_7_ = 0.031 (0.0001,10)/day, *ψ*_3_ = 0.073 (0.040,0.118), *ψ*_5_ = 0.016 (0.003,0.260), *ψ*_6_ = 0.223 (0.124,0.260), *ψ*_7_ = 0.247 (0.067,0.400), *α*_1_ = 10.120 (3.686,16.275), *α*_2_ = 0.814 (0, 20).

Both probabilistic models assume that the dynamics of coinfection within the leaf depends on the product of frequency of cells infected with either of two viral variants and the frequency of uninfected cells in the leaf (eqn. (25)). We found that removing *S_k_* term in these models resulted in significantly poorer fit of the data (results not shown). Intuitively, the frequency of uninfected cells drives the dynamics of infection and when *S_k_* approaches 0, infection of the leaf mostly stops, thus, over-predicting the data. However, when such a term is absent in equation for coinfected cells *M_k_*, co-infection would proceed even when single infections stop. With this mechanistic/mathematical insight it was difficult to come up with a biological explanation for why coinfections are dependent on the frequency of uninfected cells. One possibility that infections stop not because the number of uninfected cells declines to zero, but because of leaf-specific immune response makes uninfected cells in the leaf resistant to infection – similar to the alternative model 9 for the dynamics of infected/uninfected cells that we considered earlier (eqn. (17)).

We found interesting that the model in which the frequency of coinfected cells due to within-leaf dynamics grows logistically (eqns. (26)–(27)) could not well describe the data (Table 3). The model underestimated the frequency of coinfected cells at early time points (results not shown). This result argues that the high odds ratio for the coinfection of cells observed in our data is not likely to arise due to adjacent cells being coinfected with the two TEV variants at once. This model prediction can be tested experimentally, for example, by using microscopy and examining spatial distribution of foci of cells infected with individual viral variants or with both variants.

#### Dynamics of coinfected cells compared to singly-infected cells

While our analysis provided solid evidence that coinfection of plant cells by two TEV variants does not proceed randomly we sought to investigate how coinfection rate varies with the frequency of single-infected cells. Previous mathematical modeling-based work on HIV infection of target cells suggested that the frequency of doubly-infected cells should scale as square of the frequency of single-infected cells [42]. As far as we are aware such prediction has not been tested for plant-infecting viruses. For every leaf we therefore plotted the relationship between the frequency of coinfected cells versus the frequency of cells infected with Venus (Figure 7A-D) or BFP (Figure 7E-H) strains of TEV and compared these data with predictions of the two alternative probabilistic models. We also fitted a line to log-log transformed frequencies and estimated the slope n of the relationship (Figure 7). Several interesting results emerged.

**Figure 7:**
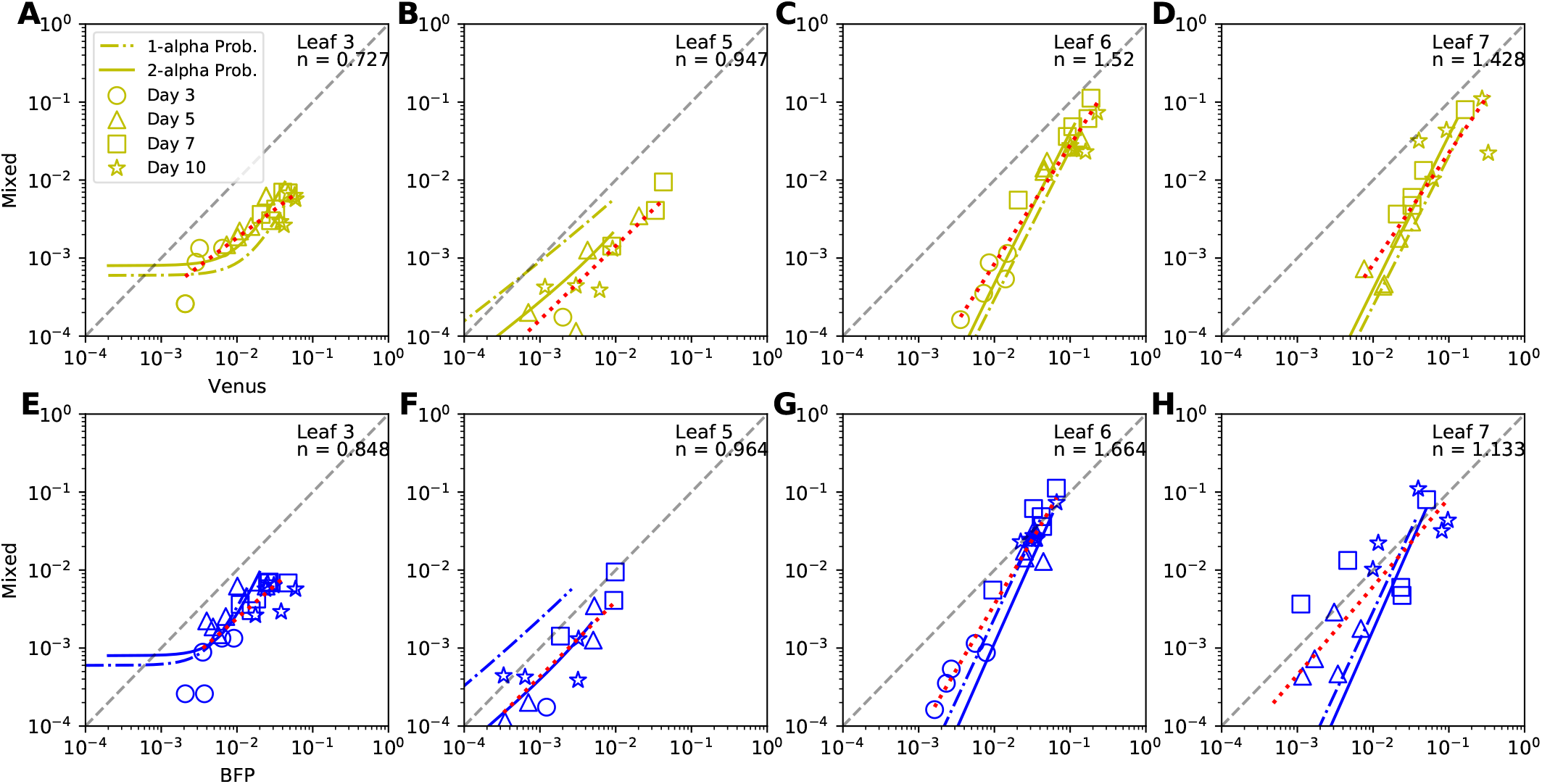
The two-alpha probabilistic model accurately describes relationship between frequency of coinfected cells and those infected with a single virus for most data. We plot the relationship between frequency of cells coinfected with Venus and BFP strains of TEV and the frequency of cells infected with a single strain (A-D for Venus and E-H for BFP) for various leaves of the plant (different panels) and different days since infection (shown by markers). Some of the data are not shown on log-log plot due to zeros of the number of infected or coinfected cells. Solid lines are the predictions of 2-alpha probabilistic model (eqn. (25)) and dash-dotted lines are the predictions of 1-alpha probabilistic model (eqn. (24)). The red dotted lines are power functions fitted to the data. The exponents of these functions are shown in the top right corners below the leaf number. Asymptotic relationship between the frequency of coinfected and single infected cells appear to become a straight line for all three models, a feature which we examine in eqns. (S.41)–(S.48). We also provide a python-based script in which these data and model predictions can be explored further (Figure S5).

First, we found that the relationship between frequency of coinfected cells and cells infected with a single virus is either sub-linear or linear for lower leaves (leaves 3 and 5, respectively, Figure 7A-B and E-F). This is not fully consistent with the results found using odds ration (Figure 5A-B) suggesting that different ways of data analysis may result in different conclusions. However, for upper leaves we found strong deviation from the linear relationship whereby coinfection frequency increased more rapidly than linearly with increasing frequency of single-infected cells (*n* > 1, Figure 7C-D and G-H). This is consistent with what we found using odds ratio (Figure 5C-D). Predictions of our best fit 2-alpha probability model were mostly consistent with the data except for the leaves 6-7 and cells, singly infected in BFP variant (Figure 7G-H). Finally, we noticed that at later time points (~ 7-10 days post infections), all of the curves in Figure 7 approximate lines. To understand why this occurs, and what the slopes of these lines are, we performed additional analyses (shown in Supplemental Information).

## Discussion

In this paper we performed extensive analyses of the recently published data on the kinetics of infection of *N. tabacum* with two variants of TEV [1]. We found that the pathway of virus dissemination in the plant could not be robustly determined directly from the data on the change in frequency of infected cells in different leaves over time — several alternative models that assumed slightly different pathways of dissemination fitted the data with very similar quality. The model assuming that viral dissemination starts from the upper leaf 7, however, fitted the data poorer than the models assuming that dissemination starts with lower (3^rd^) leaf suggesting that these data do contain some information on the direction of virus dissemination.

The best performing model in our analysis was dependent on the method of how the models were fitted to data; fitting the models using binomial distribution-based likelihood (eqn. (29)) suggested that alternative model 2 (eqn. (6)) was the best (Table S1). On the other hand, when total infection models were fitted using the least squares method based on log-transformed data and model predictions (eqn. (35)), Tromas *et al*. [1] model and alternative models 1&2 (eqn. (4) and eqn. (6)) fitted the data with the best quality (Table 2). The way experimental measurement errors influence the data remains poorly understood, and therefore, which statistical model – log-transformed least squares or binomial distribution-based likelihood – are more appropriate in fitting the models to such data remains undefined. The way forward is to better understand sources of errors in experiments measuring the fraction of infected protoplasts by flow cytometry.

It is generally unknown why not all cells in the leaves were infected by 10 days post infection; for example, leaf 3 had less than 10% of its cells infected by the end of experiment (Figure 3). One possibility that not enough time has passed for all cells to be infected. Tracking virus infection at longer than 5.5 week periods may be complicated because at this time plant physiology changes dramatically due to development of flowers. Tromas *et al*. [1] assumed that infection stops after the fraction of infected cells in a leaf exceeds some critical value, *ψ_k_*. We showed that an alternative model in which infection of a given leaf slows down due to a time-dependent factor and not directly due to increase in the fraction of infected cells, can describe the data with similar quality (Table S2). Such time-dependent factor may be an immune response such as RNAi generation and dissemination via plasmodesmata that may render cells in the leaf resistant to infection. Another factor could be changes to plasmodesmata themselves, like the accumulation of callose at the pores, that prevent the local cell-to-cell movement of the virus [43]. Physiological changes in the leaves in a growing plant may also contribute to the increased resistance of some plant cells to infection.

Our main findings, however, are about coinfection of cells with two different variants of TEV. Interestingly, by using odd ratio metrics [30] we found significantly higher frequency of coinfections of leaf cells by two viruses, in some cases with *OR* = 100 or more that is much higher than that observed in other systems [30], and in contrast with another study finding suppression of coinfections [44]. Importantly, we developed a series of novel mathematical models that track the coinfection dynamics; the best fit model also predicted higher rates of coinfections of plant cells with two viruses for the within-leaf virus spread but not for virus dissemination to other leaves (Figure 6). Additional analysis showed that at least for upper leaves (leaf 6&7) the frequency of coinfected cells increases more rapidly than linear with frequency of single-infected cells (Figure 7), and we show analytically that this is not expected in the random infection model. It has been proposed that deviation of coinfection frequency from random is likely to result from heterogeneity in target cell susceptibility to infection [30]. However, given the mechanics of virus spread in plants via plasmodesmata, ability of multiple viruses to enter the same cell, and thus, increase chances of coinfection, remains a possibility (although the model assuming this mechanism did not fit the data with best quality, (Table 3))[45, 46]. Given that virus coinfection of leaf cells in other systems can be high and that virus coinfections may result in higher virus production by infected cells [47, 48], impact of coinfections on virus evolution has received considerable attention [49–51].

As far as we are aware, Tromas *et al*. [1] performed the first comprehensive analyses of virus dissemination in plants, and so far, no similar works (experiments and modeling) on virus dissemination within and between multiple tissues have not been performed in animals. However, several studies have investigated how, for example, hepatitis C virus (HCV) spreads locally in the liver [52–54]. There is also evidence for local spread of influence A virus in humans and animals (reviewed in [55]), and mathematical models that take into account physiology of the lung tissue to study virus spread have been proposed [56]. Our observation of potential cooperativity between viruses infecting individual cells extends the results found with animal viruses such as HIV or vaccinia virus [30, 42, 57, 58]. Our analyses thus illustrate that additional insights can be generated by experiments in which infection accumulation (and loss) are tracked over time systematically in the whole organism; using barcoded viruses may be particularly useful in this regard [59].

Our study has several limitations. In our analysis we ignored the complexity of the growing, 4 week old *N. tabacum* plants, and changes that occur with leaves in the growing plant. Plants do not have pumping systems like animals, and therefore systemic movement of viruses must follow the already established pathways provided by the phloem, typically, from source to sink tissues including leaves. However, it is not always obvious based on visual appearance when a given leaf changes from being a sink to being a source (or vice versa). Viruses can manipulate source-sink relationships in their hosts; e.g., some viruses can convert source tissues into sinks [26, 60]. While we had information on the fraction of infected cells in different leaves, spatial aspects of the infection process were lost during protoplast extraction. Better understanding of virus dissemination kinetics is likely to benefit when such spatial details are also recorded, along with the high throughput flow cytometry-based measurements.

While we provided evidence that coinfection occurs at higher frequency than predicted by the random infection hypothesis, we were unable to provide a solid explanation for this effect. Both variability in susceptibility of cells to viral infection and local, cell-to-cell virus transmission via plasmodesmata may be contributing. We also have not addressed interactions between the different viral strains and the possible outcomes of these interactions.

We showed that inference of the best fit model depends on the method used to fit models to data. Given limited understanding the sources of errors in these data, the most appropriate statistical models that take into account measurement errors will need to be developed. In particular, fitting the models to data using binomial distribution-based likelihood found large differences in quality of how alternative models fitted the data. We hypothesize that binomial distribution-based likelihood amplifies small differences in the infection rate of individual leaves at early time points, leading to significant favoring of one model over the other. However, this method does not truly account for experimental noise in extraction efficiency of protoplasts from the leaves and false positives when detecting fluorescence signal from individual cells by flow cytometry. Therefore, we believe that the finding that there is one best fit model among the alternative models when models are fitted using binomial distribution-based likelihood is insufficient to choose a specific model. Additional experiments that better address experimental errors in measuring the fraction of infected cells in different leaves will be needed to derive a better statistical model to fit our dynamical models to such data.

Similarly to Tromas *et al*. [1] we ignored the fact that infection occurs in a plant, and pooled all data together without tracking infection per plant. It is clear, however, that some plants may have more infection in all leaves than others (e.g., Figure S1) and fitting the models to such “paired” data may provide additional insights into details of viral spread locally and systemically. Finally, we showed that the pathway of virus dissemination in plants cannot be easily determined from experiments that measured virus accumulation in different leaves over time, although this result was dependent on the way the models were fitted to data.

Biases introduced by extraction of protoplasts for use with flow cytometry, remain unclear. For example, infected cells may preferentially die during the extraction process which would reduce the fraction of infected cells measured. One potential way to understand such biases coming from immunology has been comparing the flow cytometry-based measurements with microscopy-based measurements [61, 62].

Our study opens avenues for future research. In particular, similar analyses may need to be performed for other plant viruses. TEV is a potyvirus, one of the largest classes of viruses in plants, and understanding dissemination of smaller viruses may prove to be interesting [63]. Together with the geminiviruses, they are responsible for the majority of disease in commercial agriculture, and understanding how these viruses disseminate in their hosts, may bring practical benefits (e.g., [64]). To better understand details of local virus dissemination it will be necessary to combine measurements of virus spread in individual leaves with flow cytometry-based measurements of the fraction of infected cells. Local virus spread can be measured by confocal microscopy and larger spread by light microscopy [24, 44, 65, 66]; previous studies have developed frameworks of how such local viral spread may be modelled [54]. Future studies should better understand why infection of a given leaf stops when not all cells are infected. Whether this is related to changes in leaf physiology (moving from sink to source) or immune responses in the leaf or systemically needs to be tested in experiments and modelled appropriately. Whether measurement of infection in leaves is sufficient to accurately predict virus dissemination kinetics is unclear. For example, roots are typical sink tissues in plants and it is likely that virus accumulation in the roots precedes or coincides with systemic virus dissemination to upper above-ground structures. Future experiments and modeling studies may benefit to include the dynamics of virus-infected cells in the plant roots. Finally, more precise understanding of the pathway of virus dissemination will benefit from additional data in which infection is initiated in different leaves and experiments in which some leaves are removed after a specific time period (e.g., [25]). Such experiments are not without caveats because removing a leaf may induce some systemic changes in the plant that in turn may influence virus replication kinetics. Therefore, such experiments would be helped by mathematical models which can make quantitative predictions on the impact of different leaf removal on the virus dissemination kinetics, and these models can be tested and some falsified [34]. Ultimately, a combination of well designed experiments to test specific hypotheses and quantitative mathematical models is likely to bring novel insights into how viruses disseminate in their plant hosts.

## Abbreviations

ODE: ordinary differential equations
TEV: Tobacco etch virus
N. tabacum: Nicotiana tabacum
*nll*: negative log-likelihood
LS: least squares
LOD: limit of detection
AIC: Akaike Information Criterion
SSR: sum of squared residuals
OR: odds ratio

## Data sources

The data for our analyses have been provided by S. Elena. Formatted data are available with this publication as a supplement (csv file) and via Github (https://github.com/Plant-Virus-Spread/Models-And-Tools.git).

## Codes

We performed most of our analyses in python and provide several key codes to ensure reproducibility of our results on github (https://github.com/Plant-Virus-Spread/Models-And-Tools.git). Specifically, we provide the codes to 1) plot the predictions of Tromas *et al*. [1] model with their parameter estimates, 2) fitting [1] model to the data using either binomial distribution-based likelihood or least squares, 3) fitting the 2-alpha probabilistic model to the coinfection data using binomial distribution-based likelihood, and 4) sliders code allowing to explore the relationship between coinfected and singly-infected cells (in the data and predictions of 2 alpha probabilistic model).

## Author’s contributions

VVG had the initial idea of the study and obtained data from the published study. VVG and JM developed the mathematical models. JM performed all the data analyses, mathematical model development and fitting to data. VVG verified some of the results in R. TBS contributed biological insights into viral spread in plants and critical overview of the modeling results. JH wrote first draft of the paper which was sequentially edited by VVG and JM. All authors contributed to writing the final version of the paper.

## Acknowledgements

We would like to thank Mark Zwart and Santiago Elena for sharing data from their published work with us. We also would like to thank members of Tessa Burch-Smith lab for comments on earlier versions of this paper. This work was in part supported by the undergraduate research award to JM and VVG, and in part by NIH (R01 GM118553) award to VVG.

## Supplemental Information

### Additional figures and tables

**Figure S1:**
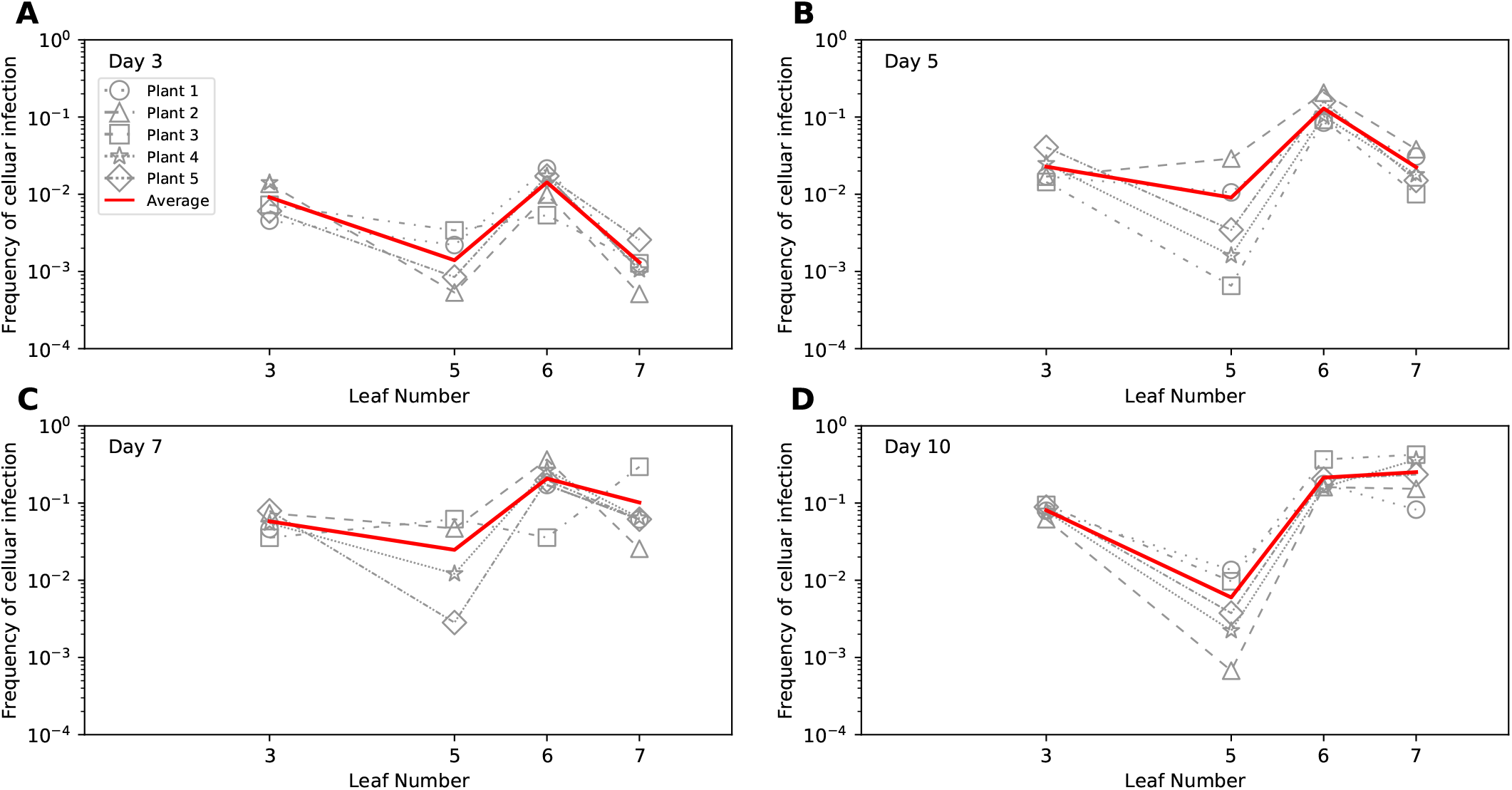
Kinetics of TEV dissemination in *N. tabacum* plants partitioned per individual plant show variability in leaf infection levels. We plot the data on infection of cells with either or both variants of TEV for individual leafs of a given plant for 3 (A), 5 (B), 7 (C), or 10 (D) days since infection. Symbols denote the frequency of infected cells in a leaf with lines connecting measurements in individual plants. Solid red line denotes average infection per leaf for a given time point.

**Figure S2:**
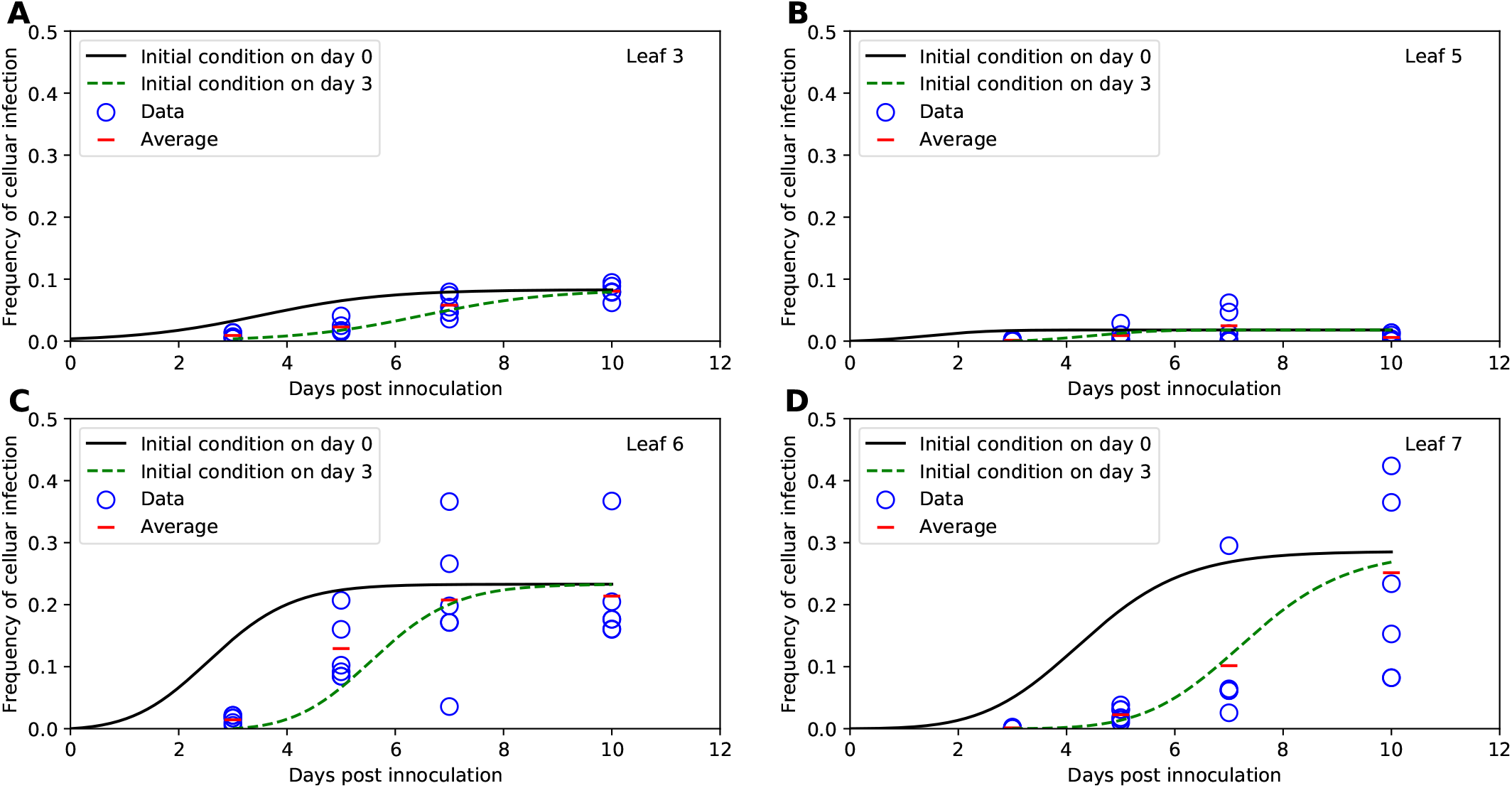
Shifting original model predictions (found in Tromas *et al*. [1]) by three days reasonably well matches experimental data. We integrated the original model (given in eqns. (1)–(3)) using an ODE solver in python either assuming that infection starts at day 0 (solid lines) or infection starts at day 3 (dashed lines); data are shown by markers for leaf 3 (A), leaf 5 (B), leaf 6 (C), and leaf 7 (D). By default, ODE solver in python is initialized by the first time point provided in the data which is day 3 in the data set. We overrode the default by forcing the solver to start infection at day 0.

**Figure S3:**
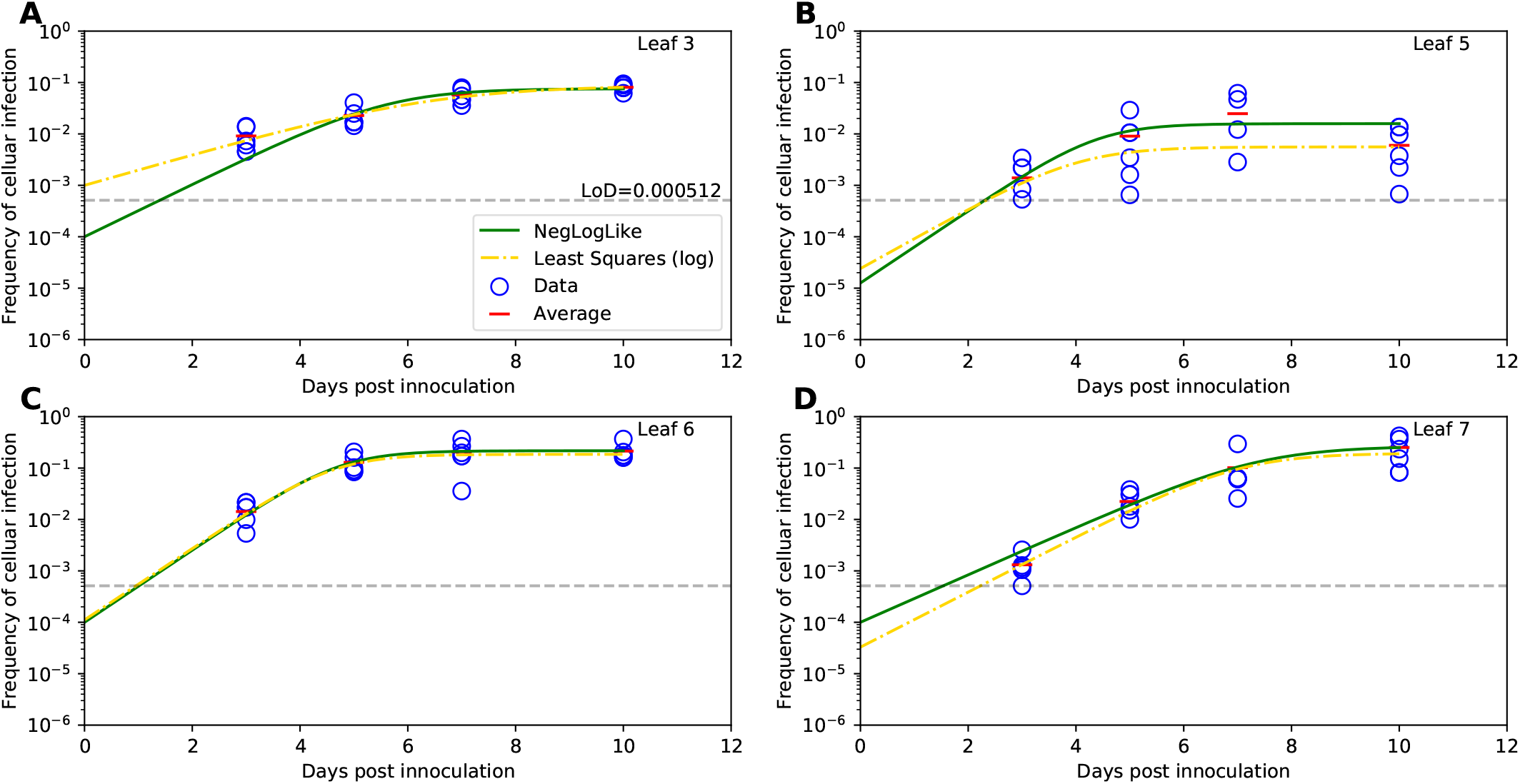
A logistic model for virus spread assuming that infection of each leaf starts and proceeds independently of other leaves fits the data with similar quality as the original virus dissemination model. We fitted the logistic model (eqn. (16)) to the virus spread data using least squares with a logarithmic transform (eqn. (35)) and assuming that infection of all leaves starts at day 0 and proceeds independently of other leaves. Data are shown by markers, solid black lines are the fits of the Tromas *et al*. [1] model, and dashed orange lines are fits of the logistic model. Parameters values and 95% confidence intervals are: *I*_0_3__ = 0.001 (0.0004, 0.003), *I*_0_5__ = 0.00002 (1 · 10^−7^, 0.001), *I*_0_6__ = 0.0001 (0.00001, 0.0007), *I*_0_7__ = 0.00003 (2 · 10^−6^, 0.00009), *β*_3_ = 0.695 (0.455, 0.895)/day, *β*_5_ = 1.348 (0.182, 3.000)/day, *β*_6_ = 1.605 (1.100, 2.194)/day, *β*_7_ = 1.236 (1.010,1.965)/day, *ψ*_3_ = 0.087 (0.084, 0.132), *ψ*_5_ = 0.006 (0.004,1.000), *ψ*_6_ = 0.185 (0.151, 0.226), *ψ*_7_ = 0.194 (0.079,0.373). The *SSR_Log_* value for this model fit was 52.285 giving an *AIC_SSR_Log__* of −10 (rounded to nearest whole number), which compared to the best model (eqn. (4)) gives Δ*_AIC_* = 6. Horizontal dashed lines in the panels denote the limit of detection LOD = 5.12 × 10^−4^.

**Table S1:** Alternative models for viral dissemination fitted to data using binomial distribution-based likelihood describe the data with different quality based on AIC values. We performed the same analysis as Table 2 except that models were fitted to data using binomial distribution-based likelihood.

| Fitted with binomial distribution-based likelihood method |  |  |  |  |  |  |  |  |
| --- | --- | --- | --- | --- | --- | --- | --- | --- |
| Parameter | Original | Alt.<br>Model 1 | Alt.<br>Model 2 | Alt.<br>Model 3 | Alt.<br>Model 4 | Alt.<br>Model 5 | Alt.<br>Model 6 | Alt.<br>Model 7 |
| $I_0$ | .0003 | .0005 | .0004 | .00006 | .0001 | .0006 | .0001 | .0001 |
| $\beta$, 1/day | .950 | .837 | .887 | 1.289 | 1.040 | 1.000 | 1.037 | 1.032 |
| $\chi_3$, 1/day | N/A | N/A | N/A | N/A | .391 | .042 | .385 | .064 |
| $\chi_5$, 1/day | .167 | .120 | .135 | .775 | N/A | 1.876 | .239 | .047 |
| $\chi_6$, 1/day | 1.046 | 5.489 | 4.964 | 8.101 | 2.275 | N/A | 2.269 | 2.251 |
| $\chi_7$, 1/day | .029 | .059 | .465 | .972 | .193 | .022 | N/A | N/A |
| $\psi_3$ | .080 | .083 | .081 | .075 | .074 | .075 | .072 | .072 |
| $\psi_5$ | .016 | .017 | .016 | .016 | .016 | .011 | .016 | .016 |
| $\psi_6$ | .224 | .223 | .223 | .217 | .225 | .235 | .219 | .219 |
| $\psi_7$ | .269 | .276 | .276 | .590 | .265 | .268 | .269 | .269 |
| $nll$ | 378317 | 378212 | 378172 | 378343 | 378495 | 379377 | 378442 | 378524 |
| $AIC_{Lik}$ | 756652 | 756442 | 756362 | 756704 | 757008 | 758772 | 756902 | 757066 |
| $\Delta_{AIC}$ | 290 | 80 | 0 | 342 | 646 | 2410 | 540 | 704 |

**Table S2:**
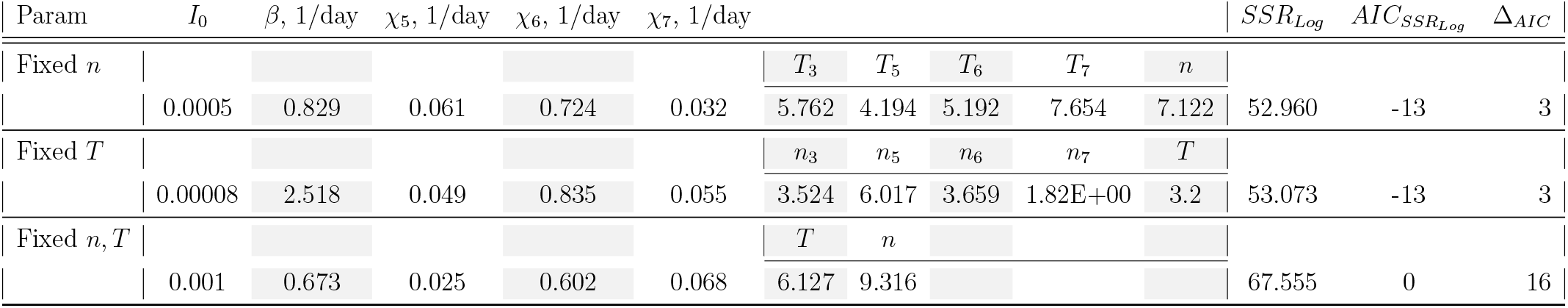
A mathematical model assuming that virus dissemination is influenced by the leaf-specific and systemic immunity can describe the experimental data. We changed the original, Tromas *et al*. [1] model by assuming the time-dependent and leaf-dependent *S_k_* function (see eqn. (17)) and fitted the model to the data using least squares with a logarithmic transform eqn. (35). In fits we either varied the time (*T_k_*) at which *S_k_* declines to zero, the Hill coefficient (*n_k_*) which determines the speed at which *S_k_* declines to zero, or both parameters being independent of the leaf number (*k*). The resulting *SSR_Log_* and *AIC_SSR_Log__* values for different model fits are shown (AICs are rounded to the nearest whole number).

**Figure S4:**
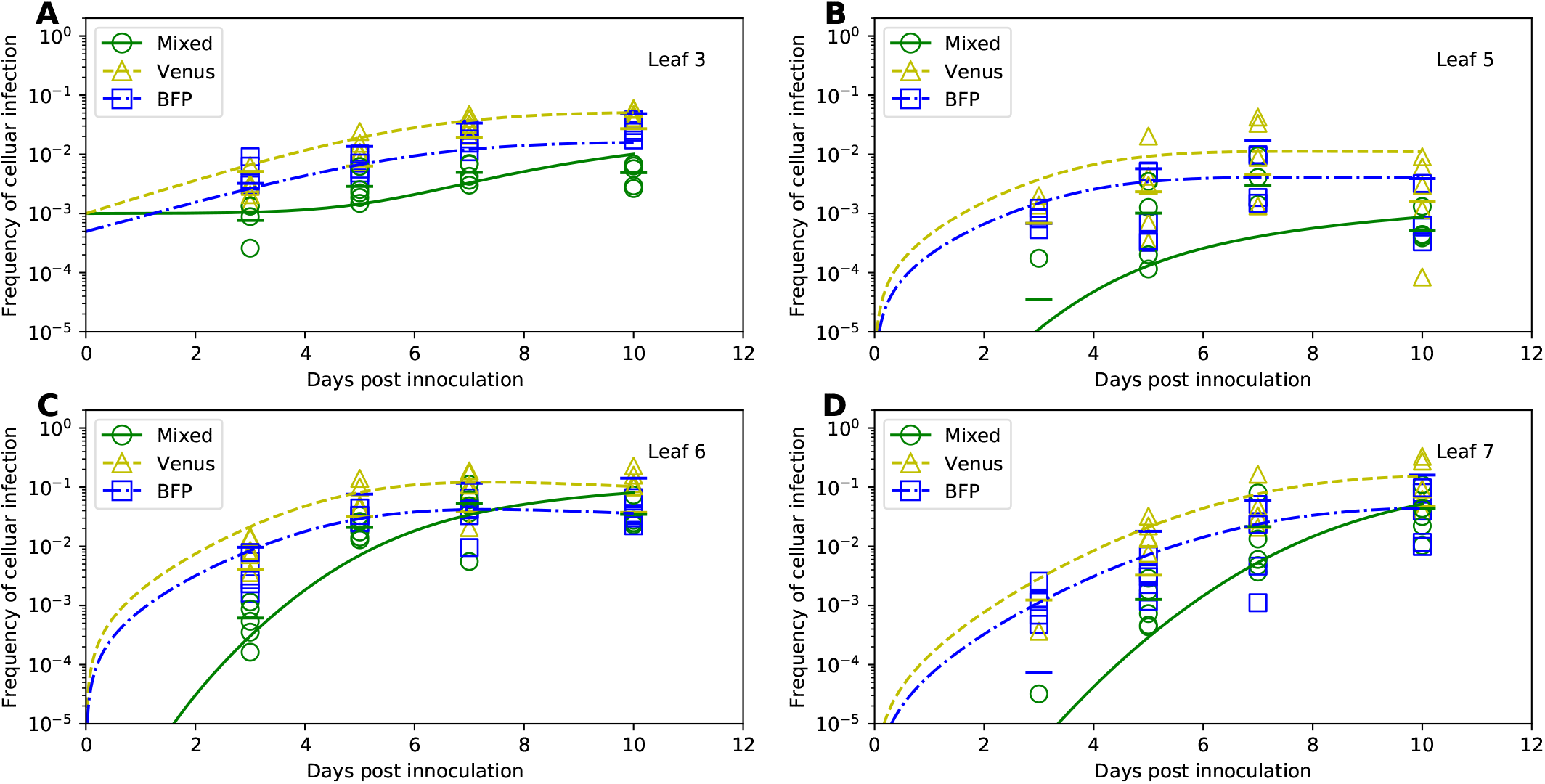
The simplest 1-alpha coinfection model does not adequately describe coinfeciton data. The 1-alpha coinfection model (given by eqns. (18)–(19) and eqn. (21)) assumes that coinfection of individual cells by two different strains occurs independently (α = 1) or coinfection may be more (*α* > 1) or less (0 < *α* < 1) likely that infection of an uninfected cell (Figure 2A). Other graph details are similar to those given in Figure 6. The parameters and 95% confidence intervals for this model are: *V*_0_ = .0006 (.0003, .001), *B*_0_ = .0003 (0.3 × 10^−5^,.001), *M*_0_ = .0001 (.0004,.001), *β_V_* = .744 (.443, 6.178)/day, *β_B_* = .666 (.260, 6.580)/day, *χ*_5_ = .269 (.034, 8.76)/day, *χ*_6_ = .939 (.395, 2.292)/day, *χ*_7_ = .044 (.015, 2.107)/day, *ψ*_3_ = .078 (.044,.954), *ψ*_5_ = .016 (.006, .028), *ψ*_6_ = .201 (.156, .267), *ψ*_7_ = .254 (.100, .449), *α* = 2.332 (.103, 5.772).

**Figure S5:**
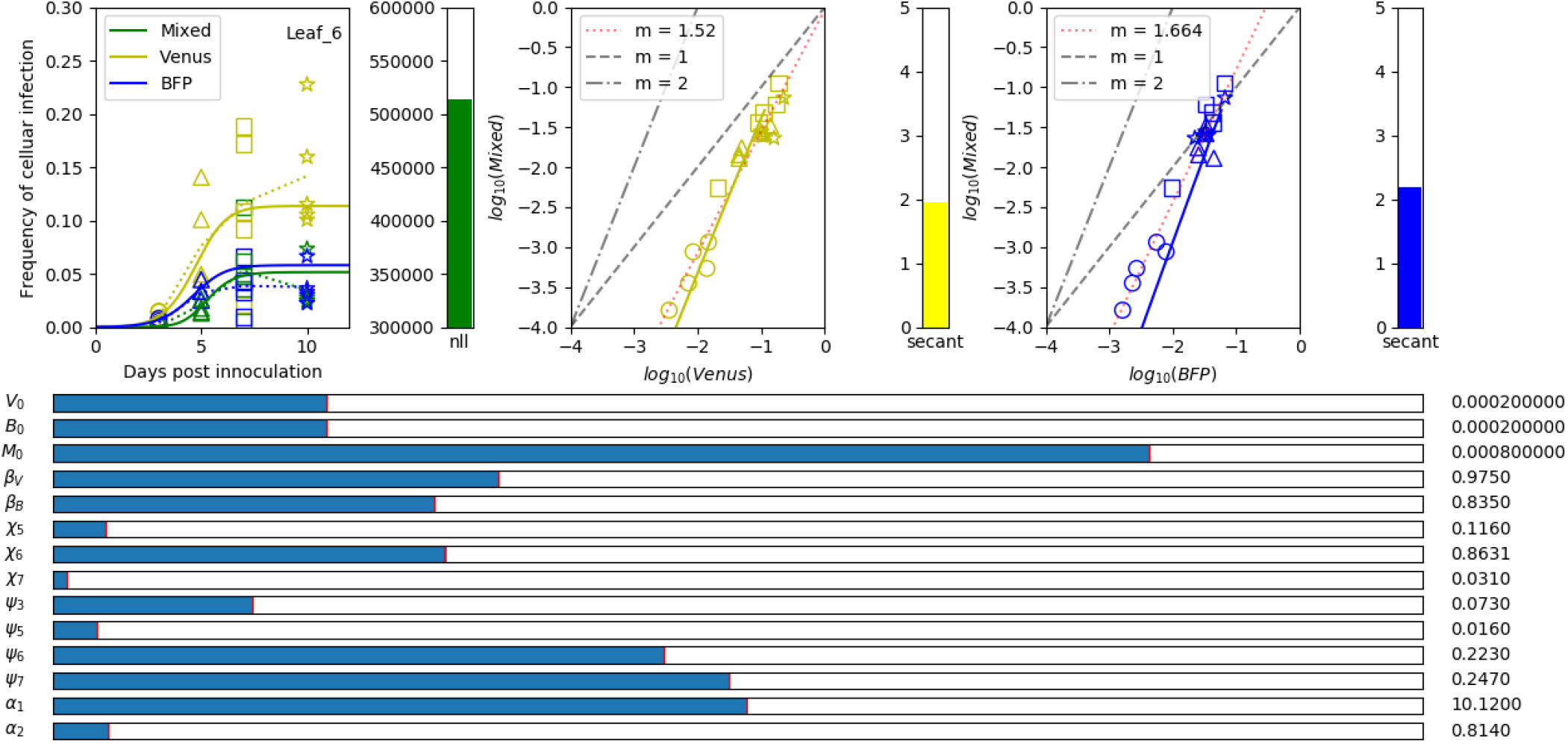
A python-based tool to study impact of parameters of a given mathematical model on the infection rate of a given leaf. By using a function Slider from pylab in python we visualized the dynamics of cell infection by individual viruses (Venus and BFP) and coinfection of cells by the two viruses according to the Probabilistic Analytic Model. Parameters of the model can be changed using sliders resulting in the changed kinetics of virus infection (shown in the left panel), or changes in the predicted relationships between the degree of coinfection of cells by two viruses (denoted as “Mixed”) and singly infected cells (Venus or BFP for middle and right panels, respectively). The example shown is for infection of leaf 6; the code allows to chose any individual leaf for visualization. In all panels data are shown by markers and predictions of the model by lines. Additional parameters shown are i) the negative log-likelihood (nll, see eqns. (30)–(33)); ii) the average ratio of the frequency of coinfected cells to singly-infected cells (secant); iii) the values of the expressions 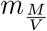 and 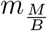 evaluated at the rightmost timepoint in the leftmost panel, in this case *t*=12 (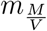 and 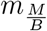.

### Expansion of the Tromas *et al*. [1] model

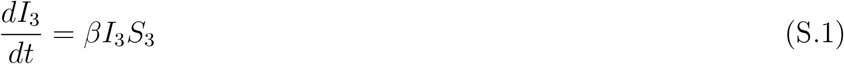

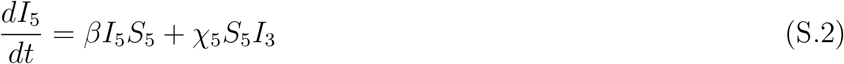

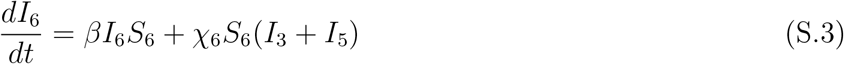

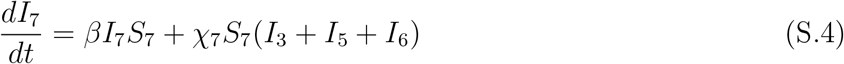

### Alternative formulations of coinfection models

Expansion of the 1-alpha model:

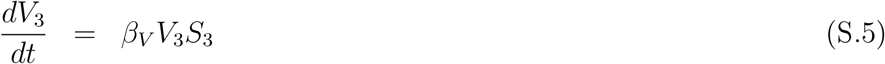

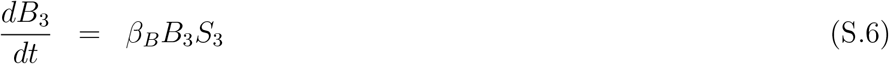

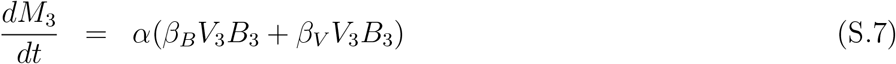

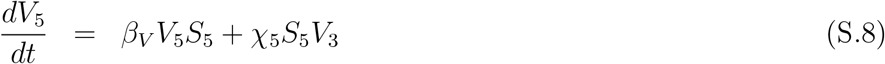

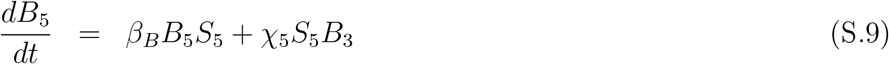

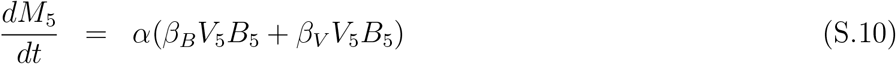

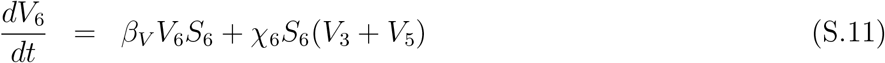

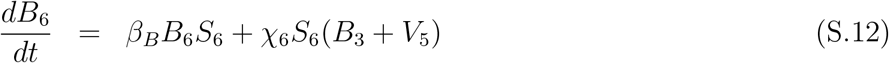

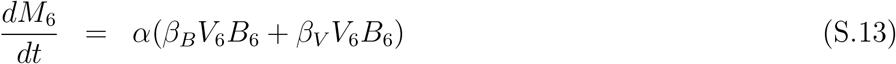

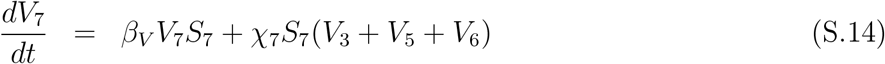

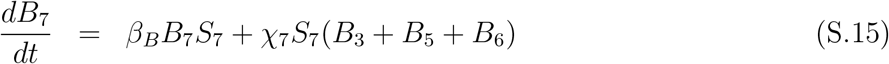

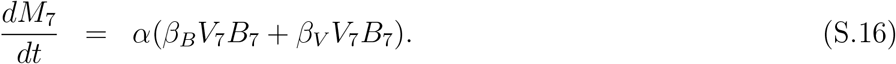

Expansion of the 2-alpha model:

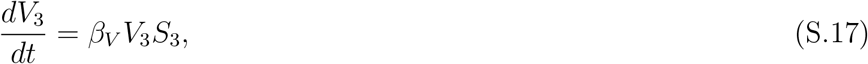

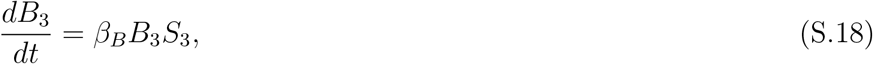

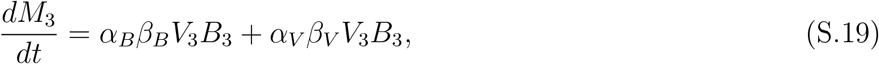

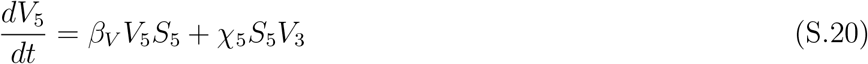

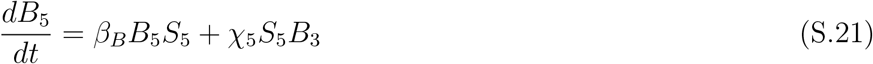

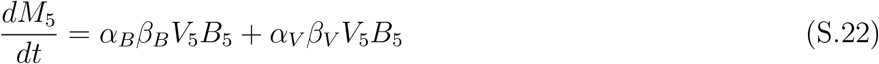

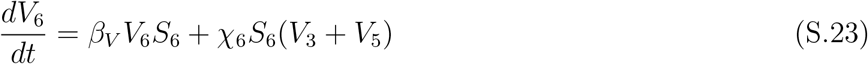

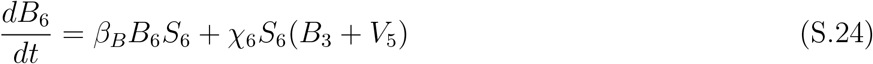

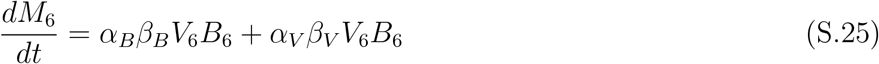

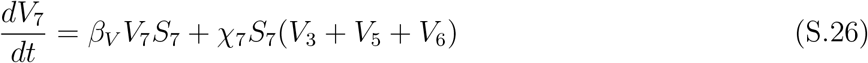

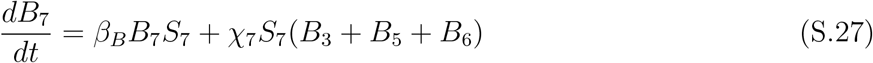

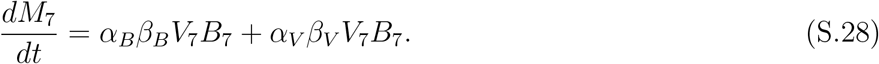

Expansion of the 1-alpha probabilistic model

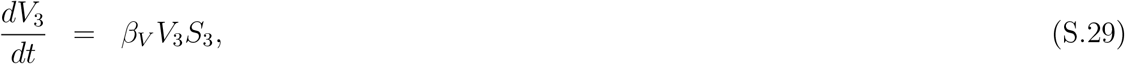

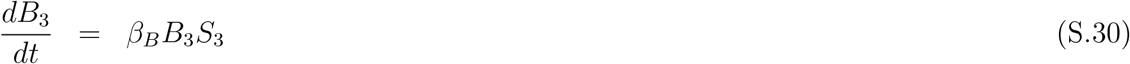

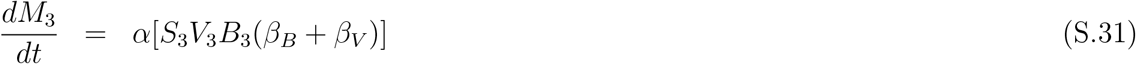

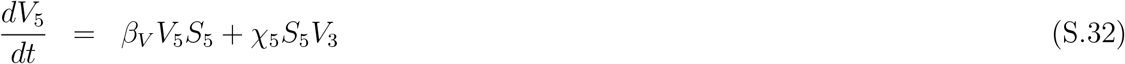

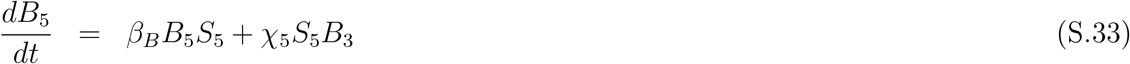

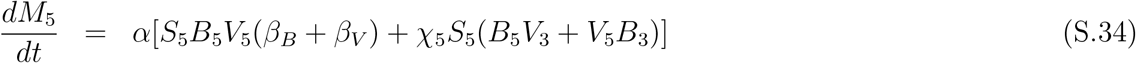

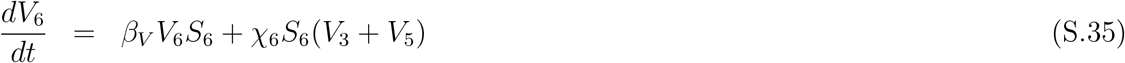

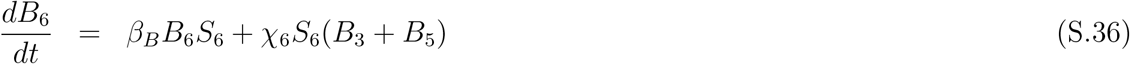

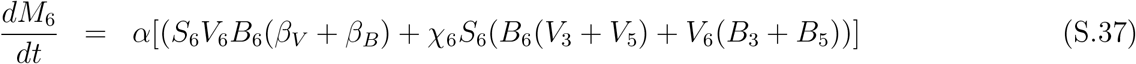

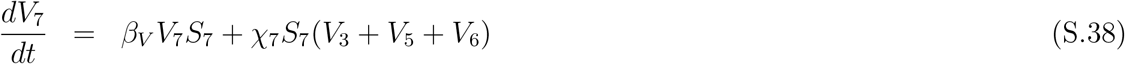

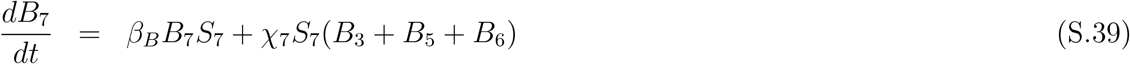

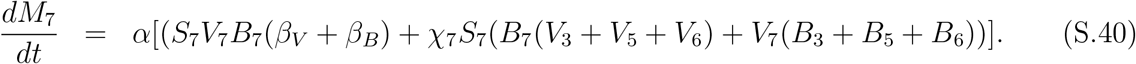

### Deriving relationship between coinfected and single-infected cells

We found that the relationship between the frequency of coinfected cells and of singly infected cells is approximately linear (e.g., Figure 7). To understand this we performed the following analyses. Specifically, we aim at calculating asymptotic behavior of 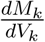 and 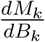.

#### Derivation of the “*V_k_*” case

Using basic calculus and eqn. (18), eqn. (19), and eqn. (23) we find:

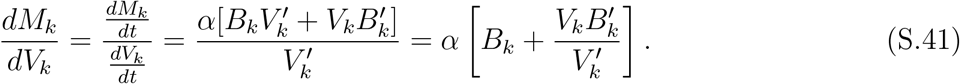

where ‘denotes derivative in time. The key to the behavior of the relationship between coinfected and single-infected cells thus lies in understanding the behavior of

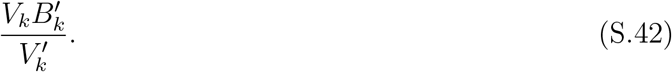

##### Leaf 3

We first consider the leaf 3 as it is the simplest and provides a method we can use to understand patterns for higher leaves. Simplifying eqn. (S.42) gives:

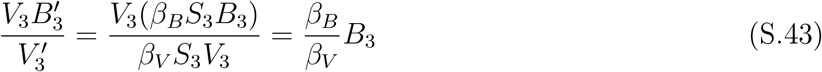

Using this, we can find the expression for the original equation.

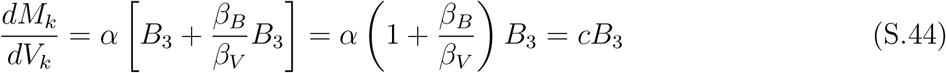

##### Further leaves

In the cases where *k* > 3 we have:

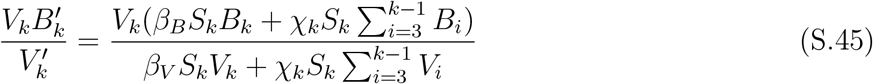

This expression is much more difficult to simply than the *k* = 3 case. However, if we take a linear combination between V_3_, V_5_, etc. we can proceed. For simplicity, we can use the average:

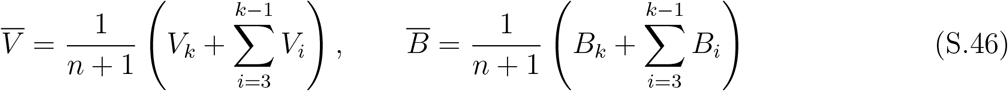

where *n* is the number of proper leaves below the *k*th leaf. Using this eqn. (S.45) becomes:

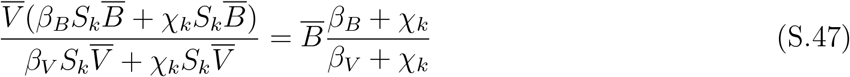

And thus we have:

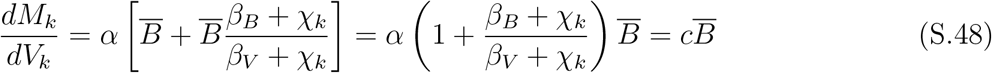

Because 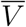 and 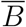 are linear functions of *V_k_* and *B_k_*, we can conclude that indeed 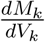 is proportional to *B_k_*, and by inference, 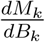 is proportional to *V_k_*.

#### Derivation of the “*B_k_*” case

Proceeding similarly as with eqn. (S.41) we find

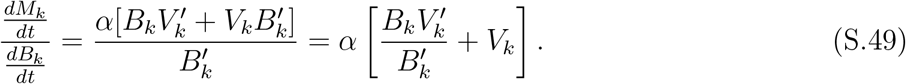

##### Leaf 3

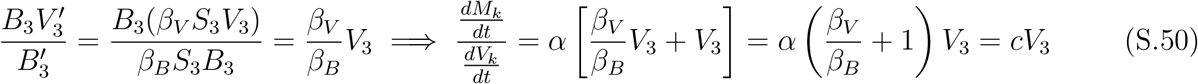

##### Further leaves

In the cases where *k* > 3 we have:

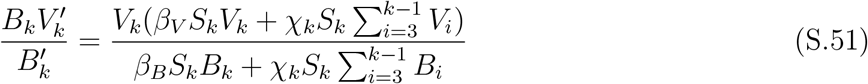

Let

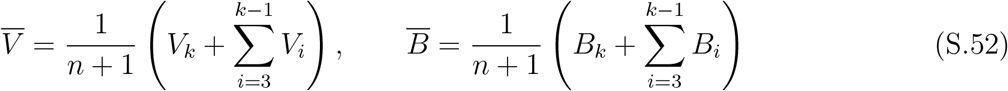

where *n* is the number of proper leaves below the *k*th leaf.

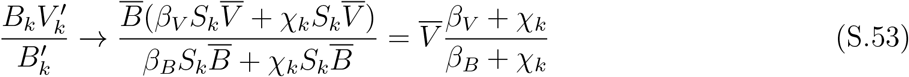

And thus we have:

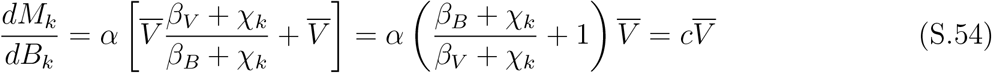

